# Cross-tissue multiomics reveals that *Akkermansia muciniphila* counteracts metabolic syndrome by reprograming gut microbiota, oleoylethanolamide and the gut-hypothalamus axis

**DOI:** 10.1101/2022.09.06.506855

**Authors:** Sung Min Ha, In Sook Ahn, Thomas Kowal, Justin Yoon, Christine A. Olson, Graciel Diamante, Ingrid Cely, Guanglin Zhang, Susanna Sue-Ming Wang, Karen Garcia, Zhejian Zhang, Angelus Cabanayan, Ruoshui Liu, Elaine Y. Hsiao, Xia Yang

**Author notes:** Correspondence: Dr. Xia Yang, Ph.D., Department of Integrative Biology and Physiology and Department of Molecular and Medical, Pharmacology, UCLA. These authors contributed equally (Sung Min Ha, In Sook Ahn).

## Abstract

High fructose diet is a major risk factor for metabolic syndrome (MetS). The gut bacterium *Akkermansia muciniphila* (*A. muciniphila*) has been shown to improve fructose-induced MetS, but the underlying mechanism remains unclear. Here, we investigated how *A. muciniphila* modulates fructose-induced MetS using multitissue, multiomics studies encompassing gut microbiota, plasma and gut metabolome, and hypothalamus single cell RNA-sequencing. *A. muciniphila* colonization enriched beneficial gut bacteria, increased metabolites including bile acids, endocannabinoids, and vitamins, and activated genes related to oxytocin and vasopressin signaling in hypothalamic neurons. Multiomics network analysis prioritized the metabolite oleoylethanolamide (OEA), an endocannabinoid analogue, as a potential regulator of gut-hypothalamic interaction conferred by *A. muciniphila,* its associated beneficial bacteria, and bile acid remodeling. Oral administration of OEA to fructose-fed mice recapitulated *A. muciniphila* effects, including counteracting body weight gain, enhancing thermogenesis, and ameliorating glucose intolerance. Concomitantly, OEA supplementation stimulated expression of its receptors and tight junction genes in the intestine, as well as neuronal activation marker *c-Fos* and oxytocin and vasopressin signaling genes in the hypothalamus. These findings underscore the regulatory role of *A. muciniphila* in gut microbiota homeostasis and metabolomic reprogramming, and pinpoint OEA as a key mediator of its action on the gut-hypothalamus axis in alleviating fructose-induced MetS.

## Introduction

Metabolic syndrome (MetS), characterized by three or more of conditions including hyperlipidemia, hyperglycemia, hypertension, insulin resistance, and abdominal fat accumulation, is a risk factor for type 2 diabetes, cancer, metabolically-associated fatty liver disease, and cardiovascular disease in humans and animal models ^1–5^. Unhealthy diets including high fat western diets and high fructose diets are main contributors to MetS. The mechanisms underlying MetS induced by different risk diets, especially the less studied fructose, is critical to counteract the MetS epidemic and develop precision medicine.

Recent studies of high fructose-induced MetS have revealed multiple mechanisms. First, high fructose reprograms the metabolic, epigenomic, and transcriptomic profiles of the host, especially in tissues such as the hypothalamus, liver, and adipose tissue ^6–9^. Fructose also interacts extensively with the gut to affect disease risk as fructose alters the gut microbiota and its metabolic outputs, and modulates gut permeability ^10^. Additionally fructose is first absorbed and metabolized in the small intestine; high fructose consumption overwhelms intestinal fructose metabolism and absorption, leading to fructose spillover to the liver and colon ^11^. Conversely, host genetics and gut microbiome composition influence fructose metabolism as well as susceptibility to fructose-induced metabolic diseases ^8,11–13^. For example, C57BL/6J (B6) mice harbor high *Akkermansia muciniphila* (*A. muciniphila*) abundance and strong resistance to obesity and glucose intolerance, whereas DBA mice, which have markedly low levels of this bacterium, have high susceptibility to those phenotypes ^12^. Treatment of susceptible DBA mice with *A. muciniphila* rendered mice resistant to fructose-induced obesity and glucose intolerance, supporting *A. muciniphila* as a potential therapeutic probiotic against fructose-induced MetS ^12^. Additionally, we found *A. muciniphila* abundance was uniquely correlated with the expression of energy homeostasis-related genes such as *Oxt*, which encodes oxytocin, in the hypothalamus ^12^, suggesting communication between *A. muciniphila* and the hypothalamus in the context of fructose consumption. However, how *A. muciniphila* confers gut-brain crosstalk to mitigate MetS remains unresolved.

We carried out a cross-tissue multiomics study that utilized single cell RNA-sequencing (scRNA-seq) of the hypothalamus, 16S rRNA gene sequencing of the fecal microbiota, and global metabolomics of fecal and plasma samples, to dissect the molecular cascades involved in the protective effect of *A. muciniphila* (Figure 1A). Our integrative multiomics analysis reveals metabolite-mediated communication from gut *A. muciniphila* to the host hypothalamus to regulate energy and metabolic homeostasis that is unique to high fructose-induced MetS.

**Figure 1.**
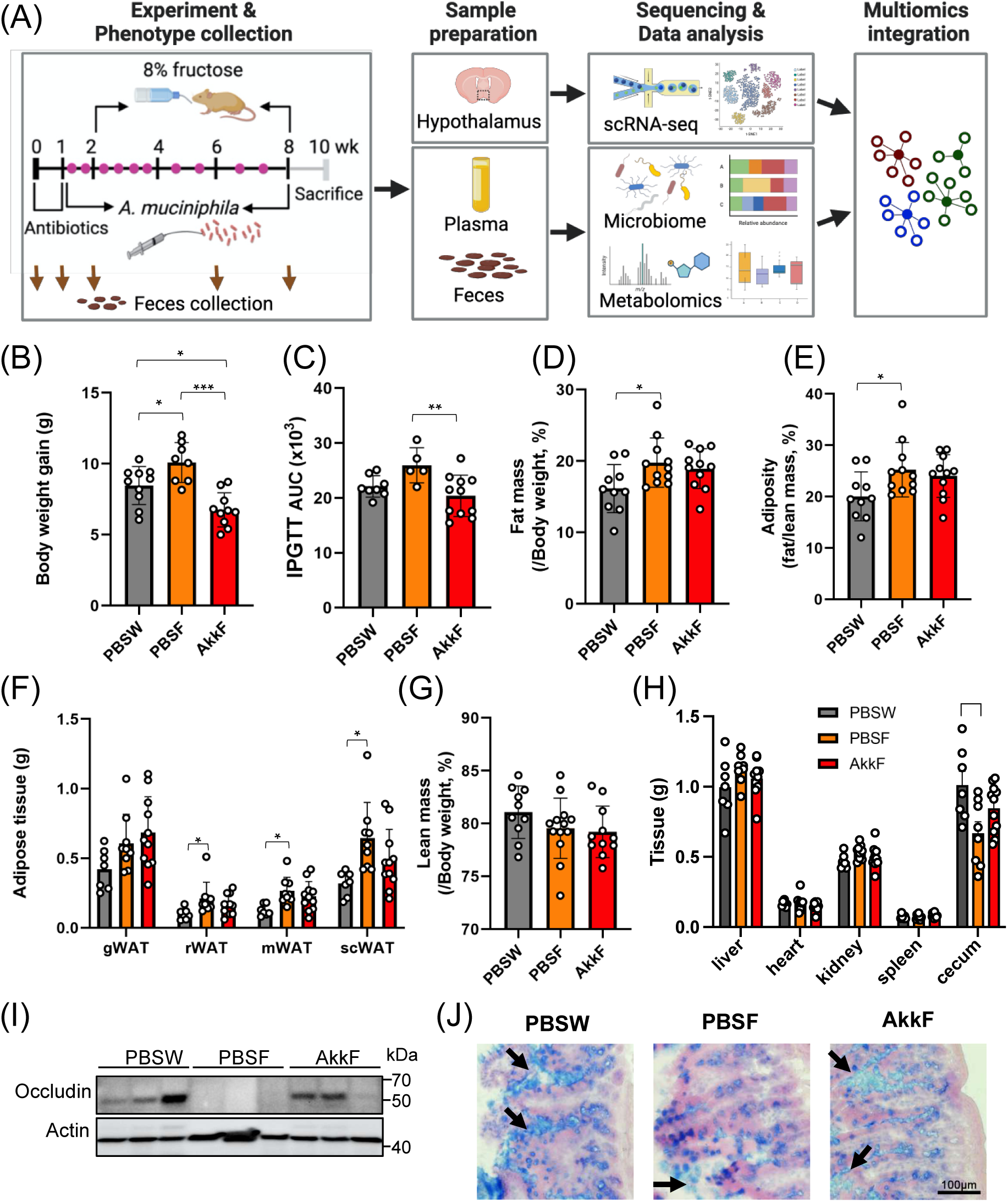
*A. mucinipihla* treatment ameliorates obesity caused by fructose diet. (A) Overall study design. DBA mice were treated with *A. muciniphila* by oral gavage (pink dots) twice weekly for 3 weeks, followed by once weekly, under 8% fructose administration. Feces were collected at week 0, 1, 2, 6, and 8. In the end of experiment, phenotypes of body weight, adiposity, and glucose tolerance were measured, followed by tissue collection between week 8 and 10. The data from microbiome, metabolomics, and scRNA-seq was generated from feces, plasma and feces, and hypothalamus, respectively, and analyzed at individual omics layer. Using individual omics data, multi-omics integrative analysis was performed to visualize systematic view of host-microbe interaction. (B) Body weight gain. (C) Glucose tolerance accessed by IPGTT, quantified as area under the curve (AUC). (D-H) Body composition changes in DBA mice after 8 weeks of fructose with or without *A. muciniphila* treatment, with (D) fat mass, (E) adiposity, (F) individual adipose tissue masses, (G) lean mass, and (H) organ weights. gWAT, gonadal white adipose tissue; rWAT, retroperitoneal white adipose tissue; mWAT, mesenteric white adipose tissue; scWAT, subcutaneous white adipose tissue. (I) Western blot analysis of tight junction related protein occludin. Actin was used as a loading control. (J) Alcian blue staining of mucin in goblet cell and between villi also confirms mucin layer was restored after *A. muciniphila* treatment. Arrows indicate mucin layer between villi. Data are presented as means ± SEMs, n = 5-13/group. Statistical significance was determined by One-way ANOVA with Tukey’s post hoc test. P-values were considered statistically significant at * p-val < 0.05, ** p-val < 0.01, *** p-val < 0.001.

## Materials and methods

### Experimental Model and Subject Details

Five-week-old DBA/2J (DBA) male mice (18-23g) were obtained from the Jackson Laboratory (Bar Harber, ME, USA) and housed in a pathogen-free barrier facility at the University of California, Los Angeles. Mice were fed chow diet (Lab Rodent Diet 5001, LabDiet, St. Louis, MO). DBA mice were chosen due to their high susceptibility to fructose-induced obesity and their lack of *A. muciniphila* based on previous studies ^8,12^.

### Animals and study design for fructose and *A. muciniphila* treatment

After one-week acclimation, 6-week old mice (baseline, week 0) were subjected to antibiotics (Ab) with a solution of vancomycin (50 mg/kg, Sigma, Saint Louis, MO), neomycin (100 mg/kg, Sigma, Saint Louis, MO), and metronidazole (100 mg/kg, Sigma, Saint Louis, MO) twice daily (9 a.m. and 6 p.m.) for 7 days according to methods previously described before weekly gavage with *A. muciniphila* (ATCC BAA-835) ^14^. Ampicillin (1 mg/mL) was also provided *ad libitum* in drinking water. Antibiotic-treated mice were maintained in sterile cages with sterile food and water and handled aseptically for the remainder of the experiments. Animals showing signs of illness after antibiotics treatment were excluded from the study. Antibiotic-treated mice were randomly divided into four groups (n=8-14/group): *A. muciniphila* gavage with regular water consumption (AkkW), *A. muciniphila* gavage along with *ad libitum* consumption of 8% fructose (NOW Real Food, Bloomingdale, IL) dissolved in water (weight/volume) starting at one week after the initiation of *A. muciniphila* gavage for 8 weeks (AkkF), PBS gavage (as control for *A. muciniphila*) with regular water (PBSW), PBS gavage with 8% fructose consumption starting at one-week after the initiation of PBS gavage for 8 weeks (PBSF). For *A. muciniphila* treatment, 200 μL *A. muciniphila* suspension (5 x 10^9^ cfu/mL in pre-reduced PBS) was gavaged throughout the experiment as previously described^15^.

Metabolic phenotypes, plasma, feces, and hypothalamic samples were collected from DBA mice using the protocols previously described ^12^. In the Ahn et al. paper ^12^, we have reported body weight gain and glucose level during intraperitoneal glucose tolerance test (IPGTT) in *A. muciniphila*-treated DBA mice. On top of these, additional phenotypic traits such as body composition (fat mass, lean mass), adiposity, and tissue weights (adipose, liver, heart, kidney, spleen) were showcased in this study. Body mass composition was determined by NMR in a Bruker minispec series mq10 machine (Bruker BioSpin, Freemont, CA). We monitored daily food and water intake as well as a total calorie intake. Fecal samples were collected at weeks 0 (beginning of Ab treatment), 1 (beginning of *A. muciniphila* treatment), 2 (beginning of fructose treatment), and 6 at the beginning of the 12-h dark cycle (6 p.m.). Mice were fasted overnight before sacrifice, Blood samples were collected from the retro-orbital sinus at the end of fructose feeding (week 10 from baseline) after a 12-h overnight fast and transferred to blood collection tubes with K2 EDTA (BD, Becton Dickinson, USA). Plasma was obtained by centrifugation for 10 minutes at 1,500 x g. After blood collection, mice were sacrificed, and hypothalamus samples were dissected as described in the previous study ^16^. All samples were snap frozen and stored at -80°C until further analysis. Plasma samples were used for metabolomics; fecal samples were used for 16S rRNA sequencing and metabolomics; hypothalamic samples were used for scRNA-seq.

Animal studies were performed in accordance with the National Institutes of Health Guide for the Care and Use of Laboratory Animals. All experimental protocols were approved by the Institutional Animal Care and Use Committee at the University of California, Los Angeles.

### 16S rRNA sequencing of fecal samples

Total bacterial DNA was isolated from fecal samples using ZymoBIOMICS DNA Miniprep extraction kit (Zymo Research, USA) according to manufacturer’s instructions. Bacterial 16S rRNA was amplified for amplicons spanning the variable region 4 (V4) using barcoded primers. The amplicons from different samples were pooled, purified, and sequenced using the Illumina MiSeq4000 instrument with average sequencing depth of 10,531 reads per sample. Sample processing and sequencing were performed at the Microbiome Core Lab at UCLA.

### Global metabolomic profiling of plasma and fecal samples

Plasma and fecal samples at week 10 were extracted in methanol and divided into 4 aliquots: two for analysis by two separate reverse phase (RP)/UPLC-MS/MS methods with positive ion mode electrospray ionization (ESI), one for analysis by RP/UPLC-MS/MS with negative ion mode ESI, one for analysis by hydrophilic interaction chromatography (HILIC)/UPLC-MS/MS with negative ion mode ESI. Non-targeted global metabolomic profiling in plasma and feces was performed at Metabolon, Inc. (Durham, NC, USA) ^17–19^.

### scRNA-seq of hypothalamus samples

Freshly dissected hypothalami were dissociated with papain (Worthington, Lakewood, NJ, USA) for single cell isolation ^20^, and cells were suspended in 0.04% BSA-PBS at a final concentration of 100 cells/μl as previously described ^21^. Single cells were encapsulated in droplets together with barcoded primer beads (ChemGenes, Wilmington, MA, USA) to generate single-cell transcriptomes attached to microparticles (STAMPs), followed by generating cDNA library with the Drop-seq protocol by Macosko et al 2015 ^22^. Paired-end sequencing data was generated using Illumina NovaSeq 6000 at a sequencing depth of ∼40,000 read pairs per cell, with custom length of 26 bp for read 1, 76 bp for read 2, and 8 bp for read 1 index.

### Validation of OEA as a key metabolite regulating the *anti-obesity effect of A. muciniphila*

As OEA was found to be a consistent metabolite altered after *A. muciniphila* treatment in both fecal and plasma samples, we tested whether OEA recapitulates the anti-obesity effect of *A. muciniphila* in the fructose-induced obesity model. Male DBA mice of 8-week age were randomly assigned to 4 groups (n = 8/group); OEA gavage (50 mg/kg) along with *ad libitum* consumption of 8% fructose water (Fructose + OEA), OEA gavage with regular water (Water + OEA), Vehicle gavage with 8% fructose water (Fructose + Veh), vehicle gavage with regular water (Water + Veh). OEA (99%, Botany Bioscience, San Luis Obispo, CA, USA) was dissolved in vehicle [saline/polyethylene glycol/Tween 80 (90/5/5, v/v)] and administered to mice daily 30 min before the 12-h dark cycle (5:30 p.m.). OEA purity was analyzed using HPLC (S&N Lab, Santa Ana, CA), and chemical structure by H-1 NMR spectra was confirmed with OEA spectra reported by Wang et al.^23^ (NuMega Resonance Lab, San Diego, CA). The OEA treatment dose was determined by converting the manufacturer’s recommended human dose (200 mg per adult per day) to an equivalent mouse dose ^24^.

Food and drink intakes were monitored twice a week on a per-cage basis. Body weight was measured weekly. To test the thermogenic effects of OEA, body temperature was measured rectally with a BAT-12 Microprobe Thermometer (Physitemp, Clifton, NJ) at week 7. Temperature measurements were performed between 2 p.m.- 3 p.m. at room temperature (18-23^◦^C). At the end of the fructose and OEA treatment (week 8), an intraperitoneal glucose tolerance test (IPGTT) was performed. Briefly, animals were fasted for 12 hours prior to the IPGTT. Each mouse was intraperitoneally injected with 20% glucose in saline at 2g glucose/kg body weight. Blood glucose levels from tail vein were measured at 0, 15, 30, 90, and 120min post glucose injection using an AlphaTrak blood glucose meter (Abbott Laboratories, North Chicago, IL). Mice were fasted overnight before sacrifice, and hypothalamus and ileum samples were dissected. All samples were snap frozen and stored at -80°C until further analysis.

### Real time qRT-PCR

To test downstream targets of OEA treatment in the intestine, total RNA was extracted from ileum and hypothalamus using Direct-zol Miniprep Kit (Zymo Research, Irvine, CA) and cDNA was synthesized using cDNA Reverse Transcription Kit (Thermo Fisher Scientific, Madison, WI). Real time qRT-PCR was performed in QuantStudio 3 Real-Time PCR System (Thermo Fisher Scientific, Madison, WI) for the relative quantification of *Ppara, Gpr119, Nape-pld, Faah, or Cd36* gene expression from ileum which were previously implicated in the activities of OEA. We also quantified *Avp, Oxt, c-Fos, Fosb, Jun,* and Ganq gene expression from hypothalamus. PCR primers were designed using PrimerBank ^25^ and the Primer Blast tool available from NCBI web site. Melt curve was checked to confirm the specificity of the PCR product. Relative quantification was performed using the 2 ^ (−△△ CT) method. *Beta-actin* was used as an endogenous control gene to evaluate the target gene expression levels. All data are presented as the means ± SEM of n = 4-7/ group.

### Western blot

Ileum was homogenized in lipa buffer and protein was extracted. Protein concentration was measured using BCA protein assay (Bio-Rad, Irvine, CA, USA). Proteins (20μg per lane) were separated on 4-12% or 12% Mini-PROTEAN® TGX gels (BioRad, Hercules, CA, USA) and transferred onto a PVDF membrane (Pall, Ann Arbor, MI, USA). Membranes were blocked in 5% milk in TBS-T (TBS, 0.1% Tween 20), and incubated at 4 °C overnight with primary antibodies (actin or occludin) diluted in TBS-T with 5% milk. After washing, membrane was incubated in HRP (horseradish peroxidase)-conjugated secondary antibodies for 1 hour. Proteins were visualized using ECL reagents (BioRad, Hercules, CA, USA). Blot images were captured using ChemiDoc Image System (BioRad, Hercules, CA, USA).

### Mucin staining

Tissue sections of 6 μm thickness were prepared from frozen ileum embedded in OCT. Alcian blue staining was performed to stain mucin according to manufacturer’s protocol (Abcam, Cambridge, UK).

### Bioinformatics analysis 16S rRNA data analysis

Microbial sequence data was analyzed using QIIME2 pipeline (version 2020.8) ^26^. Raw FASTQ files demultiplexed by sample barcodes were used as input files for QIIME2. Denoising was conducted with DADA2 followed by amplicon sequence variant (ASV) analysis ^27^. The representative sequences from ASVs were targeted for taxonomy classification via naïve Bayesian classifier with Silva ver. 138 database ^28^. Alpha and Beta diversity were calculated with Shannon and weighted Unifrac values as input, respectively. Linear discriminant analysis effect size (LefSe) was used to compare the abundances between *A. muciniphila*-treated and control groups at each time point to obtain the linear discriminant analysis (LDA) score ^29^.

Representative ASVs from all samples were subject to further functional profiling using PICRUSTs2 ^30^. A series of standard workflows were conducted to place sequence into reference phylogeny to infer 16S rRNA copy number. Using phylogeny and 16S rRNA copy number, gene family profiles were predicted and pathway abundances were measured. All statistical differential abundance (ASV and pathways) were tested using Kruskal-Wallis with pairwise Wilcoxon rank sum exact test as a post hoc test. P-values were adjusted with Benjamini-Hochberg.

### Analysis of global metabolomic profiling of plasma and fecal samples

Raw data was extracted and proceeded with peak identification and quality control using Metabolon’s hardware and software^31^. Metabolites were identified by matching to the library based on authenticated standards that contains the retention time/index (RI), mass to charge ratio (*m/z*), and chromatographic data on all molecules present in the library. Metabolites were quantified using area under the curve of the peaks. All metabolomics samples were processed and analyzed (differentially abundant metabolites and pathway enrichments) as described previously ^32^. Briefly, differential metabolites analysis between AkkF and PBSF groups was performed using Welch’s Two Sample t-Test on log transformed data with q-val < 0.05 as an indication of significance with high confidence. In cases where significant metabolites at q-val < 0.05 were not enough to run downstream pathway analysis, p-val < 0.05 was used to select differential metabolites. Pathway enrichment analysis was performed by mapping metabolites to the Metabolon proprietary database. Pathway enrichment score was calculated by

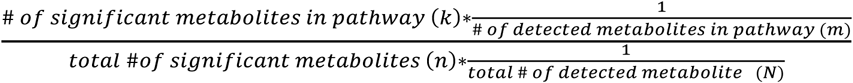

The statistical significance was calculated using a Fisher’s exact test and the top representative metabolic pathways were selected with an enrichment score greater than 1.5 and a hypergeometric p-val < 0.05.

### scRNA-seq data analysis

DropSeqPipe version 1.13 (https://github.com/Hoohm/dropSeqPipe) was used to preprocess the scRNA-seq sequencing data and generate digital gene expression matrix files. Gene expression matrix from each sample was loaded into the Seurat R package version 4.0.3. Quality control was conducted based on number of genes (200-4000), unique molecular identifiers (UMIs; 500-5000) and percent of mitochondrial genes (<10%) to filter out cells that did not meet these thresholds. Potential doublets were removed using DoubletDecon version 1.1.6 with default parameters ^33^.

After quality control, all samples were integrated together using the *FindIntegrationAnchors* function in Seurat. Next, the integrated Seurat object was used for dimension reduction using principal component analysis (PCA), k-nearest neighbors (KNN) for graph construction, and Louvain clustering to cluster cells. Cell clusters were visualized using t-distributed stochastic neighbor embedding (tSNE) and uniform manifold approximation and projection (UMAP).

Cell type identity was determined for each cluster using the reference mapping function in Seurat where hypothalamic scRNA-seq data from Romanov et al. (GSE74672) was used as a reference ^34^. The data from Romanov et al. was used as a reference because they used tissue specimen that comprises all regions in hypothalamus; paraventricular nucleus, anterior nucleus, suprachiasmatic nucleus, Dorsomedial nucleus, Ventromedial nucleus, and arcuate nucleus. Initially, the mapping resulted in seven different cell types, namely, neurons, ependymal cells, oligodendrocytes, microglia, vascular and smooth muscle (VSM) cells, and astrocytes. Of these, ependymal cells, oligodendrocytes and microglia were subject to further refinement based on cell subtype marker genes from Chen et al ^35^. Ependymal cells were divided into ependymal and tanycytes based on the expression of tanycyte specific genes (*Ccdc153* and *Rax*). Oligodendrocyte cells were separated into myelinating oligodendrocyte (MO; top marker *Mobp*), newly formed oligodendrocytes (NFO; top marker *Fyn*), and Oligodendrocyte precursor cells (OPC; top marker *Pdgfra*). Microglia was further separated into different states based on the expression of *Mrc1*, *Dab2*, and *Trem119* to represent under-differentiated microglia ^36^ (Figure S1A-C). Lastly, neuronal cells were subclustered to elucidate neuronal subtypes that express different neuropeptides. Markers for each of these sub-clusters were found using the *FindAllMarkers* function. These neuronal subclusters were then named after their corresponding uniquely expressed markers or neuropeptides.

To obtain differentially expressed genes (DEGs) resulting from *A. muciniphila* treatment for each cell type, we performed non-parametric Wilcoxon rank sum test and Poisson regression using the Seurat package for general cell types and neuronal subtypes, respectively.

Poisson test was used to identify DEGs of neuronal subtypes because the assumption of Poisson model fits better with the data within each neuronal subtype where homogenous cells expressing mRNA of a gene at fixed rate and low UMI counts (average of 1,874 UMIs/cell) ^37^. The transcriptomes of each cell cluster or subcluster were compared between groups with or without *A. muciniphila* treatment, namely AkkF versus PBSF groups, and AkkW versus PBSW groups. DEG p-values were adjusted for multiple testing using Bonferroni correction. For pathway analysis which requires a larger number of DEGs, DEGs at p-val <0.05 were used. Pathway analysis was carried out using EnrichR version 3.0, with KEGG_2019_Mouse, BioCarta_2016, Reactome_2016, and GO_Biological_Process_2018 as reference pathway databases ^38^ and significant pathways were determined using adjusted p-val < 0.05 cutoff.

### Integrative analysis across multi-omics data

Within the microbiome data, pair-wise correlation between gut bacteria was calculated using FastSpar to avoid compositional bias from relative abundances in microbiome data ^39,40^. Pair-wise correlation between *A. muciniphila*-responsive metabolites in plasma or feces and *A. muciniphila* gut microbiota, as well as between DEGs from scRNA-seq and metabolic phenotypes including body weight and weight gain were assessed using Biweight midcorrelation (bicor) ^41^. A network was generated with bacterial taxon, metabolites, DEGs, and phenotypes that had pair-wise correlations at p-val <0.05. Hub nodes were found using the weighted key driver analysis (wKDA) method in Mergeomics with default parameter ^42–44^. Subnetworks of the hub nodes were extracted using degree of 3. Network was visualized using Cytoscape (version 3.9.1) ^45^.

### Statistical analysis for metabolic phenotypes and Real-Time qRT-PCR data

Statistical significance for time-course data (body weight gain and glucose tolerance) were determined by repeated measures Two-way ANOVA, followed by Tukey’s post hoc test.

All other phenotype data were evaluated using One-way ANOVA with Tukey’s post hoc test. For analyses involving multiple variables (e.g., tissues or genes), comparisons among groups were performed separately for each variable. Statistical analyses were performed using GraphPad Prism (version 8.2; GraphPad Software, Inc.). Data are expressed as means ± SEMs. P-val < 0.05 was considered statistically significant.

## Results

### *A. muciniphila* treatment counteracts fructose-induced metabolic dysfunctions

To test the role of *A. muciniphila* in counteracting fructose-induced metabolic dysfunction, we administered *A. muciniphila* or vehicle control (PBS) with or without 8% fructose water to DBA mice for 8 weeks (Figure 1A). Prior to *A. muciniphila* administration, mice were pre-treated with broad-spectrum antibiotics for 7 days. The four treatment groups were: *A. muciniphila* + fructose water (AkkF), *A. muciniphila* + water (AkkW), PBS + fructose water (PBSF), and PBS + water (PBSW). *A. muciniphila* treatment of fructose-fed mice significantly reversed the body weight gain and glucose intolerance phenotypes (Figure 1B,C). Fructose feeding significantly increased normalized fat mass, adiposity, and adipose depot weights (PBSW vs. PBSF; Figure 1D-F). In contrast, these measures were not significantly different between PBSW and AkkF, indicating that *A. muciniphila* attenuated the fructose-induced adiposity gain, which is similar to previous findings from a high-fat diet–induced obesity model ^46,47^. Lean mass or other tissue weights were not affected by *A. muciniphila* with the exception of the cecum (Figure 1G,H). Notably, *A. muciniphila* reversed the significant reduction in cecum weight induced by fructose. Lastly, *A. muciniphila* also improved gut barrier function by restoring the fructose-induced decrease in occludin expression to normal levels (Figure 1I) and restoring the fructose-induced decrease in mucin between villi as shown by Alcian blue staining (Figure 1J). Overall, these results confirm that *A. muciniphila* ameliorates fructose-induced body weight gain and glucose intolerance while improving intestinal barrier integrity.

### *A. muciniphila* reprograms hypothalamic signaling involved in endocannabinoid–oxytocin–vasopressin axes

*A. muciniphila* has been reported to contribute to gut-brain crosstalk by modulating neurotransmitter pathways such as serotonin ^48,49^. Agreeing on the involvement of the gut-brain axis, our previous study showed that *A. muciniphila* abundance was correlated with the expression of several metabolic genes in the hypothalamus, including *Oxt, Th,* and *Bmp7* ^12^. To understand how *A. muciniphila* remodels gene expression in the hypothalamus, a key control center of whole-body metabolism, we performed hypothalamic transcriptome profiling at the single-cell level. For scRNA-seq, after pre-processing and quality control, we analyzed 4,198, 3,229, 3,046, and 3,124 hypothalamic cells from the AkkF, AkkW, PBSF, and PBSW groups, respectively. Each group consisted of three mice, for a total of 12 independent scRNA-seq samples. Among the 11 major hypothalamic cell types identified (Figure 2A,B, Figure S1,2A) ^34–36,50^, cell-type proportions were unchanged by *A. muciniphila* treatment or fructose diet for most cell types except for neurons (fructose diet effect, p = 0.009) and tanycytes (Akk x diet interaction, p = 0.012) as assessed by 2-way ANOVA (Figure 2C, Figure S2B).

To determine genes that were affected by *A. muciniphila* in each hypothalamic cell type, we analyzed differentially expressed genes (DEGs) between Akk and PBS groups on either the fructose or the control diet within each cell type. Neurons exhibited the highest DEG numbers indicating stronger transcriptional responses to *A. muciniphila*, particularly under fructose feeding with 60 DEGs at adjusted p-val < 0.05 and 646 DEGs at p < 0.05, followed by endothelial, mature oligodendrocytes (MO), astrocytes, and microglia (Figure 2D,E). *A. muciniphila*-induced DEGs in the neurons showed significant overlap between the fructose and control diet conditions (Figure 2E), but many DEGs (e.g., *Zfp91, Jund, Usp32, Jun, Mt3, Gnaq,* and *Apoe*) were found only in the fructose condition (Figure S2C, Table S1). Across cell types, DEGs such as those involved in stress responses (*Fos, Fosb, Egri, Atf3*) showed consistent direction of change under both diets, although DEGs with opposite direction of change between diets were also identified, such as *Lamp1, Zfp1, Mt1* and *Actb*.

**Figure 2.**
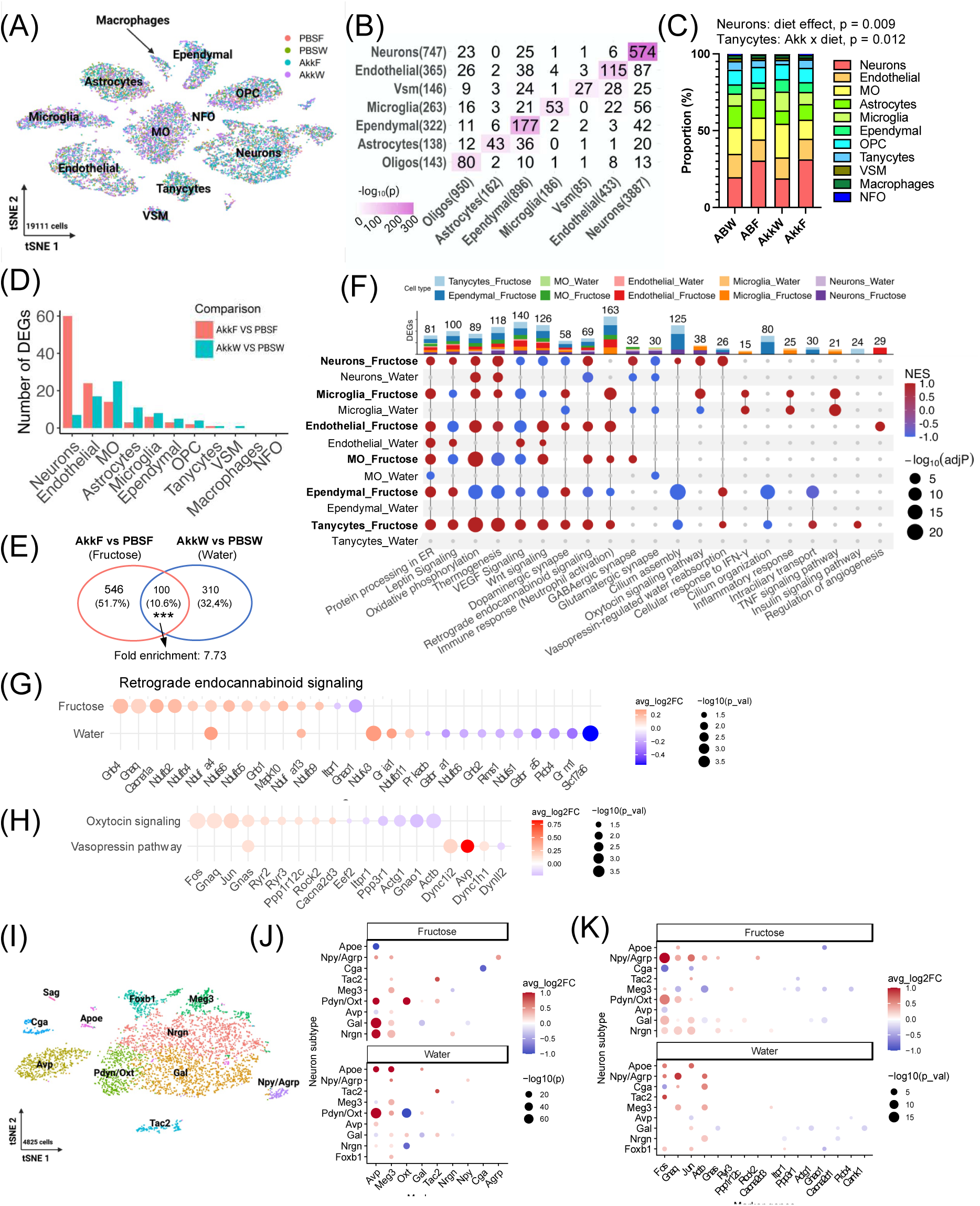
*A. muciniphila* alters the transcriptome profile of hypothalamic cell populations. (A) t-distributed stochastic neighbor embedding (tSNE) plot showing 11 different cell types. Colored dots represent cells from 4 different groups, PBSW, PBSF, AkkW, and AkkF. (B) Heatmap showing the overlap between marker genes of major cell clusters (rows) defined in the current scRNA-seq with hypothalamus cell type markers (column) from Romanov et al. ^34^. The numbers of markers for each cell type are shown in the parenthesis. The numbers of overlapping markers are shown in the heatmap cells. The statistical significance of the overlap was calculated by Fisher’s exact test and indicated by color (the darker color represents higher statistical significance). (C) Cell type proportion. (D) Numbers of DEGs (adjusted p-val < 0.05) from individual cell types affected by *A. muciniphila* under fructose or control diet condition. (E) Venn diagram showing the numbers of DEGs (p-val < 0.05) in neuronal cells. Significant overlap was determined by Fisher’s exact test. *** p-val < 0.001. (F) An UpSet plot showing enriched pathways across cell types under fructose and water conditions, with or without *A. muciniphila* supplementation. The bar plot on the top indicates the number of significantly enriched pathways unique to or shared among cell types. In the dot plot, each dot represents a pathway enriched in a given cell type and condition. The color gradient corresponds to the normalized enrichment score (NES), with red indicating upregulation and blue indicating downregulation. Dot size represents the statistical significance of enrichment (–log10 adjusted P-value). (G, H) Dot plots depicting DEGs involved in retrograde endocannabinoid signaling (G), and Oxytocin signaling pathway and vasopressin-regulated water reabsorption in neuronal cell type under fructose condition (H). (I) tSNE plot showing a total of 11 subtypes of neurons. (J, K) Dot plots showing significant expression changes of neuronal subcluster markers (J) and genes involved in oxytocin signaling (K) by A. muciniphila in fructose or water condition. Statistical significance of DEG was determined by Wilcoxon ranked sum test and poisson generalized linear model for cell types and neuronal subtypes, respectively. Sample size is n=3 independent tissues/group. DEGs were considered statistically significant at adjusted p-val < 0.05 (D) or p-val < 0.05 (E-H,J,K), with an absolute log fold change ≥ 0.1. Statistical significance of factor effects (Akk, diet, Akk x diet interaction) was determined by Two-way ANOVA (C).

Pathway analysis of the *A. muciniphila*-induced DEGs revealed protein processing in ER, leptin signaling, oxidative phosphorylation, thermogenesis, VEGF signaling, and Wnt signaling as shared across several cell types under fructose diet (Figure 2F, Table S2). In hypothalamic neurons under both fructose and control dietary conditions, mitochondrial oxidative phosphorylation and thermogenesis pathways were upregulated, suggesting *A. muciniphila* enhanced neuronal energy expenditure and metabolic signaling. Notably, although genes involved in retrograde endocannabinoid signaling were also significantly enriched in neuronal DEGs under both diets, the pathway genes showed an overall increase under fructose diet and an overall decrease under the control diet (Figure 2F, G). Retrograde endocannabinoid signaling is known to regulate presynaptic neurotransmitter release and synaptic plasticity, and to modulate neuropeptide signaling in hypothalamic circuits ^51^. Consistent with this, neuropeptide signaling pathways including oxytocin (OXT) and vasopressin (AVP) signaling, both of which regulate energy balance and metabolism ^52^, were upregulated in neurons under fructose diet only (Figure 2F, H). Among DEGs included in OXT signaling pathway, *Fos* (neuronal activation marker) expression was upregulated in both fructose and water groups, indicating that *A. muciniphila* activated neurons (Figure 2H, Figure S2D) ^53^.

To further characterize neuronal changes induced by *A. muciniphila*, we subclustered neurons into eleven subtypes based on the neuropeptide markers (Figure 2I, Figure S3A). Fructose significantly increased the proportions of Gal (p = 0.007 by Two-way ANOVA) and Npy/Agrp (p = 0.032) subtypes, whereas *A. muciniphila* induced a non-significant decrease (p=0.09) in Foxb1 neurons (Figure S3B). Significant expression changes in *Oxt*, *Avp*, *Agrp*, *Gal* and *Meg3*, regulators of hypothalamic neuroendocrine activity, energy balance, and glucose homeostasis ^54–56^ were found in the neuronal subtypes (Figure 2J). Among DEGs in the OXT signaling pathway, *Fos, Gnaq, Jun,* and *Actb* were mostly upregulated in the neuronal subtypes by *A. muciniphila*, and *Fos* was upregulated mainly in Pdyn/Oxt and Npy/Agrp subtypes; the patterns are stronger in fructose group (Figure 2K, Table S1).

Together, these results indicate that *A. muciniphila* induces broad transcriptomic changes particularly in neurons under the fructose diet condition to upregulate retrograde endocannabinoid signaling, oxytocin and vasopressin neuropeptide pathways, and thermogenic and oxidative phosphorylation gene programs.

### *A. muciniphila* treatment inhibits pathogenic microbes and promotes beneficial gut bacteria

To understand whether *A. muciniphila* drives the phenotypic and hypothalamic gene expression changes directly or via secondary shifts in the microbiota, we tracked microbial dynamics throughout the treatment process of *A. muciniphila*. *A. muciniphila* was present in most of DBA fecal samples at low levels (average 1.3%) at baseline (week 0) before antibiotic treatment (Figure S4A). After a week of antibiotic treatment, the alpha diversity (Shannon index) of all samples decreased significantly as expected (Figure 3A and Figure S4B). One week treatment with *A. muciniphila* further decreased the overall diversity at week 2. By contrast, the PBS control group showed restoration of microbial diversity at week 2 after antibiotic treatment. At week 6 (5 weeks after *A. muciniphila* treatment), *A. muciniphila* increased and became one of the dominant species in the *A. muciniphila*-treated group compared to week 2 (Figure S4C,D). Principal coordinate analysis (PCoA) plot of beta diversity (weighted UniFrac) showed samples from the *A. muciniphila* treatment group clustered apart from the PBS control samples (Figure 3B). These results support successful colonization of *A. muciniphila* by 5 weeks of treatment.

**Figure 3.**
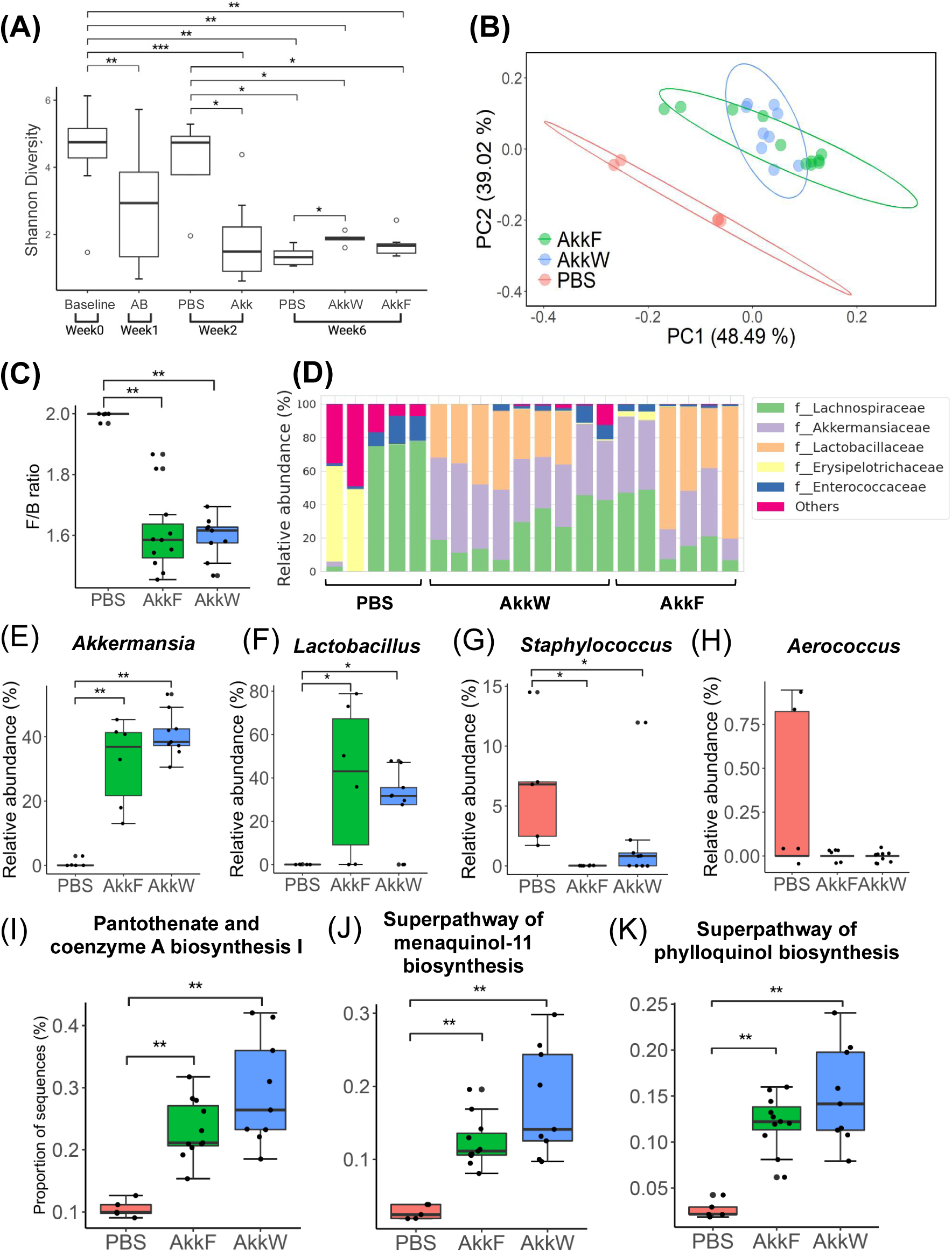
*A. muciniphila* exerts beneficial effects by inducing compositional and functional shifts of microbiota. (A) Alpha diversity (Shannon) changes after different treatment or diet was applied. (B) Beta diversity plot based on weighted UniFrac showing a clear separation between *A. muciniphila* treated groups from control (PBS) group. (C) F/B (Fermicutes/Bacteroidetes) ratio changes by *A. muciniphila* treatment. (D) Family-level microbial taxonomy structure in each sample. (E-H) Box plots showing changes of relative abundance of genera (E) *Akkermansia* (F) *Lactobacillus* (G) *Staphylococcus* (H) and *Aerococcus* by *A. muciniphila* treatment. (I-K) Prediction of functional abundance of microbiome using PICRUSt2 showing Pantothenate (Vitamin B5), menaquinol-11 (Vitamin K2), and phylloquinol (Vitamin K1) after *A. muciniphila* treatment. (B-K) Analyses were performed at week 6. Sample size is n=5-9/group. Statistical significance was determined by Kruskal-Wallis test, followed by pairwise Wilcoxon rank sum test as post hoc analyses. Values were considered statistically significant at p-val < 0.05. * p-val < 0.05, ** p-val < 0.01, *** p-val < 0.001.

At week 6, the Firmicutes/Bacteroidetes (F/B) ratio, which is positively correlated with obesity ^57^, showed a significant decrease in response to *A. muciniphila* treatment in both control and fructose diet groups (Figure 3C). Additionally, *Akkermansiaceae* and *Lactobacillaceae* are dominant families in the *A. muciniphila* treatment group (Figure 3D). At the genus level, *A. muciniphila* significantly promoted the growth of *Lactobacillus,* a well-known beneficial bacterium, while reducing *Staphylococcus* and *Aerococcus* (Figrue 3E-H). Additional analysis of all amplicon sequence variants (ASV) using the linear discriminant analysis effect size (LefSe) analysis and cladogram confirmed that genus *Akkermansia* is enriched in the *A. muciniphila* treated group, whereas in the control PBS group *Romboutsia*, *Peptostreptococcaceae*, *Aerococcus*, *Agathobacter*, and *Staphylococcus* were enriched (Figure S5A,B). Among these, *Peptostreptococcus*, *Aerococcus*, and *Staphylococcus* are known to be clinically pathogenic ^58,59^. Overall, *A. muciniphila* treatment inhibited bacterial taxa known to be pathogenic and promoted select bacterial taxa that are regarded as beneficial.

To determine the functional implications of these gut microbial changes with respect to *A. muciniphila* treatment, PICRUSt2 was used to predict the functional profiles of the bacterial community altered by *A. muciniphila* ^30^. Our results indicated upregulation of pantothenate (Vitamin B5), menaquinol (Vitamin K2), and phylloquinol (Vitamin K1) biosynthesis pathways by *A. muciniphila* treatment in both water and fructose diets (Figure 3I-K). Supplementation of all these nutrients has been reported to reduce obesity and insulin resistance ^60,61^. To the best of our knowledge, an increase in the biosynthesis of these vitamins due to *A. muciniphila* treatment has not been reported elsewhere. These observed changes in gut bacterial community and functional pathways may contribute to downstream metabolite alterations that further mediate *A. muciniphila*-host interactions.

### *A. muciniphila* induces metabolomic changes including endocannabinoids related metabolites

To determine whether *A. muciniphila* modulates metabolites that may mediate subsequent communication with host tissues, we carried out global metabolomic analysis from the fructose-consuming groups with or without *A. muciniphila* treatment. A total of 813 and 786 metabolites were identified in plasma and feces, respectively. PCA revealed a distinct separation between control (PBSF) and *A. muciniphila* (AkkF) groups for both plasma and fecal metabolite profiles (Figure 4A). Differential metabolite analysis using Welch’s Two Sample t-test identified 11 plasma metabolites and 344 fecal metabolites that were differentially abundant between AkkF and PBSF at q-val < 0.05 (Table S3 and S4). Since the small number of differential plasma metabolites identified using the stringent cutoff limited our ability to perform pathway analysis, we used a less stringent threshold (p-val < 0.05), identifying 134 plasma and 328 fecal differential metabolites for downstream analysis (Figure 4B-D).

**Figure 4.**
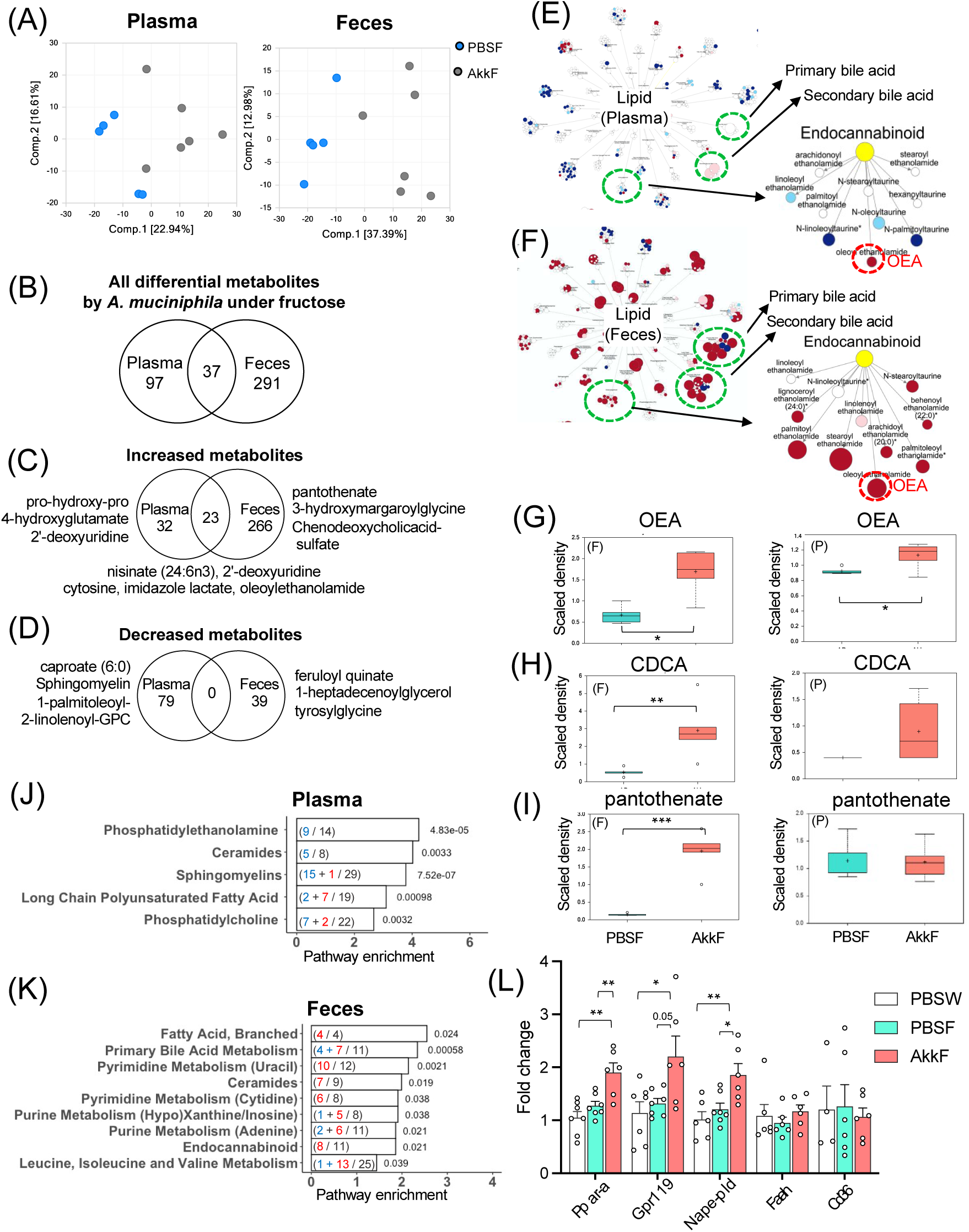
*A. muciniphila* induces metabolomic profile shift with the increase of endocannabinoid OEA. (A) Principal component analysis (PCA) plots of plasma and fecal samples assessing the variance of metabolites between *A. muciniphila* treated group and control (PBS) group under fructose treatment. The percentages of variance explained by the first (Comp. 1) and second (Comp. 2) principal components are shown in brackets. (B-D), Venn diagrams showing the numbers of (B) all differential (C) increased and (D) decreased metabolites in plasma and feces of *A. muciniphila*-treated group (AkkF) compared with PBS group (PBSF) under fructose. Differential metabolites were determined by Welch’s t-Test (p < 0.05). (E-F) Metabolites involved in lipid pathway in (E) plasma and (F) feces altered by *A. muciniphila* under fructose. Red and blue dots indicate lipid metabolites upregulated and downregulated at p-val < 0.05, pink and cyan dots indicate metabolites upregulated and downregulated at 0.05 < p-val < 0.1, respectively, by *A. muciniphila*. Dot size indicates the relative fold change. (G-I) Box plots showing the levels of (G) OEA (H) CDCA (chenodeoxycholic acid), and (I) pantothenate in feces (F, left panels) and plasma (P, right panels) between AkkF and PBSF groups. Differential metabolites between AkkF and PBSF were determined by Welch’s t-Test (q-val < 0.05). (A-I) n = 5-6/group. (J-K) Bar-plots showing top representative pathways of differential metabolites in (J) plasma and (K) feces. Enrichment hypergeometric p-values are shown beside the bars. Inside the bar, black, blue, or red indicates # of all detected, decreased, or increased metabolites in the pathway by *A. muciniphila* under fructose. (L) Fold changes of gene expression levels of *Ppara, Gpr119, Nape-pld, Faah,* and *Cd36* in the ileum. Data are presented as means ± SEMs. n = 6-7/group (except *Cd36* of PBSW, n = 4). Statistical significance of each gene was determined by One-way ANOVA with Tukey’s post hoc test and values were considered statistically significant at p-val < 0.05. * p-val < 0.05, ** p-val < 0.01, *** p-val < 0.001.

*A. muciniphila* induced stronger alterations in fecal metabolites compared with plasma metabolites (Figure S6). Metabolites in the lipid family were mostly increased by *A. muciniphila* in fecal samples but were decreased in the plasma (Figure 4E,F and Figure S6A,B), suggesting reduced lipid absorption into the circulation and increased excretion into feces. Indeed, *A. muciniphila* has been reported to reduce jejunal lipid absorption and lower serum triglyceride while increasing fecal triglycerides in HFD-fed mice ^65^. Agreeing with our functional prediction based on the microbial community changes in the previous section (Figure 3I), pantothenate (Vitamin B5) was the most significantly upregulated metabolite in feces, but not in plasma (Figure 4C and 4I).

Thirty-seven metabolites overlapped between plasma and fecal samples. These include nisinate and oleoylethanolamide (OEA) which are increased by *A. muciniphila* in both fecal and plasma samples (Figure 4C,E-G). Nisinate is a very long chain polyunsaturated omega-3 fatty acid involved in DHA biosynthesis ^62^. OEA is an endocannabinoid analogue primarily synthesized in the small intestine and a member of the N-acylethanolamine (NAE) family that acts as a neurotransmitter regulating the gut-brain axis via vagus nerve or by directly reaching the area postrema in the brainstem ^63^. OEA is known to regulate feeding, lipid metabolism, and energy homeostasis by modulating hypothalamic neuropeptide signaling ^64–66^. Other metabolites in the NAE family were also increased in feces by *A. muciniphila* treatment, including N-palmitoylethanolamine (PEA) and N-stearoylethanolamine (SEA) (Figure S7A,B). Among these NAEs, only OEA was increased in both plasma and feces (Figure 4C,E-G). Furthermore, the fecal bile acid profile was significantly altered, with increased levels of both primary bile acids such as chenodeoxycholic acid (CDCA) and secondary bile acids, notably ursodeoxycholic acid (UDCA) (Figure 4F,H and Figure S7C,D). CDCA is known to promote the activity of N-acyl phosphatidylethanolamine phospholipase D (NAPE-PLD), an enzyme that plays a crucial role in OEA biosynthesis ^67^.

Pathway analysis using MetaboAnalyst^68^ also supported the importance of OEA and the endocannabinoid pathway. *A. muciniphila* treatment significantly affected five metabolic pathways in plasma (Figure 4J), including phosphatidylethanolamine and phosphatidylcholine, both of which are OEA substrates^69^. In fecal samples, *A. muciniphila* affected nine metabolic pathways (Figure 4K) including the endocannabinoid pathway, branched fatty acids, primary bile acids, and pyrimidine metabolism. Therefore, across fecal microbiota, fecal and plasma metabolomics, and hypothalamic scRNA-seq data, the endocannabinoid pathway, known to participate in gut-brain axis signaling ^70^ was identified as a consistent feature affected by *A. muciniphila* under fructose diet; within the endocannabinoid pathway, OEA appears to be an important metabolite.

### Intestinal gene expression supports changes in OEA biosynthesis and signaling genes

As the intestine is a main site for OEA synthesis and signaling, we investigated genes involved in these pathways upon *A. muciniphila* treatment. In support of the OEA increase in our metabolomic results, real-time qPCR analysis of ileal tissues showed that *A. muciniphila* significantly increased the expression of *Napepld* (encoding enzyme NAPE-PLD responsible for OEA synthesis) without changes in *Faah* (OEA degradation) and *Cd36* (OEA substrate uptake) in fructose-fed mice (Figure 4L). OEA is a specific ligand for peroxisome proliferator-activated receptor-alpha (PPARα) and G protein-coupled receptor 119 (GPR119), both of which regulate food intake and glucose homeostasis via gut-brain signaling ^51^. Indeed, *Pparα* and *Gpr119* expression levels were significantly upregulated by *A. muciniphila* treatment (Figure 4L). These results support the notion that *A. muciniphila* treatment induces OEA biosynthesis and promotes downstream intestinal signaling genes *Pparα* and *Gpr119* to mediate communications between gut microbiota and the host.

### Multiomics integration reveals a molecular network involved in *A. muciniphila* treatment

To further elucidate the molecular cascades involved in *A. muciniphila*-to-host crosstalk based on our multiomics datasets, we performed a correlation network analysis across the microbiome, metabolome, scRNA-seq, and phenotype data (Figure 5A). Within the microbiome families, we found that *Akkermansiaceae* was positively correlated with beneficial bacteria *Lactobacillaceae, Enterobacteriaceae, Erysipelotrichaceae,* and *Clostridiaceae*, but negatively correlated with pathogenic bacteria *Staphylococcaceae*, *Peptostreptococcaceae, Aerococcaceae,* and *Streptococcaceae*. Thus, the beneficial effect of *A. muciniphila* may arise not only from itself but also from the establishment of a healthier microbial community.

**Figure 5.**
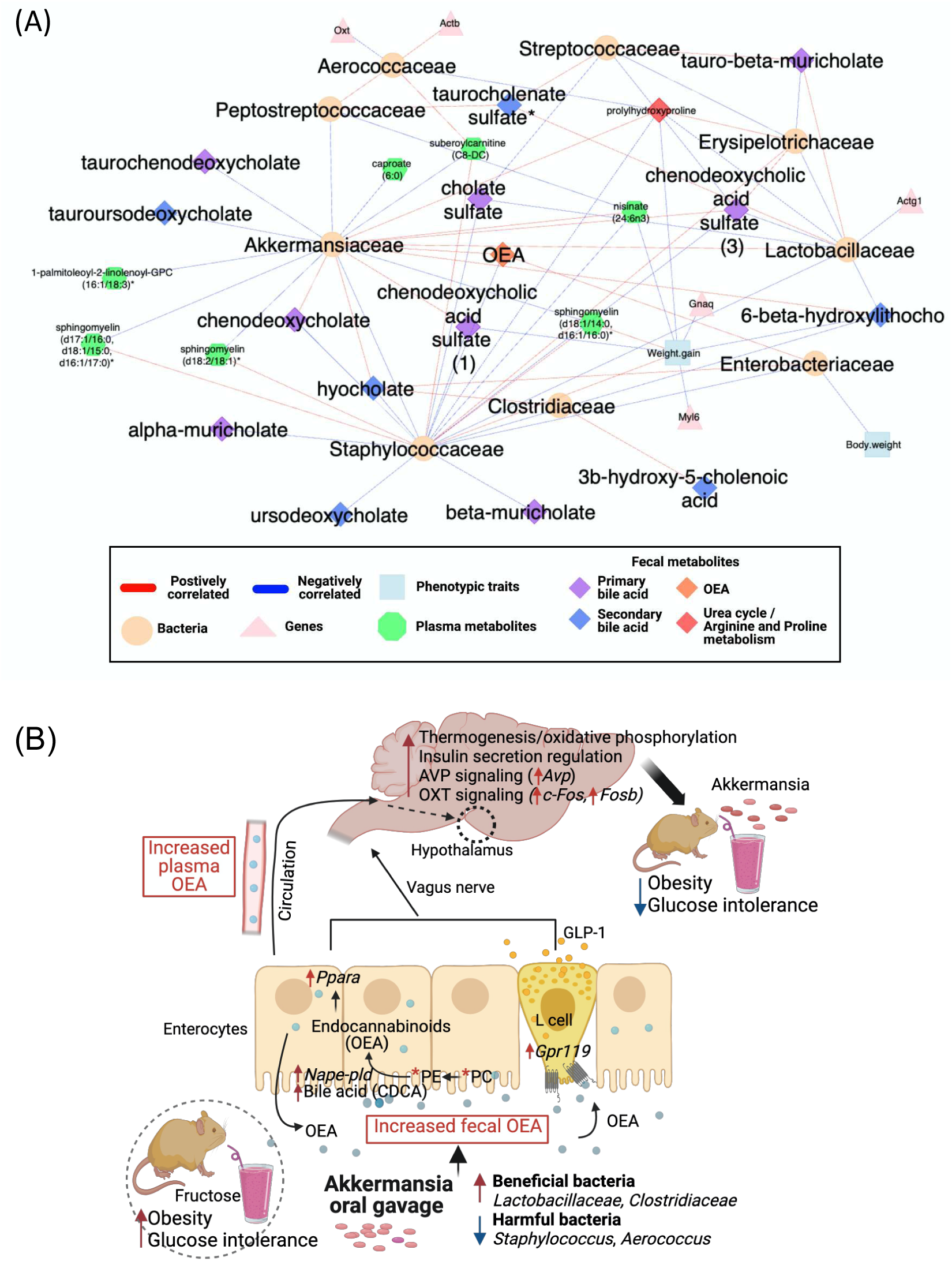
Integration of microbiome, metabolomics, and scRNA-seq data and the proposed action mechanism of *A. muciniphila* against fructose-induced obesity. (A) Correlation network between microbial taxa, fecal and plasma metabolites, DEGs from hypothalamus scRNA-seq, and metabolic phenotypes. Correlation coefficients among family-level microbiome were calculated using FastSpar (p-val < 0.05) and used as a backbone of network structure. Correlations between microbes, metabolites, genes, and phenotypes were calculated by Biweight midcorrelation (p-val < 0.05). Nodes are the bacteria and their adjacent correlated genes, metabolites, and phenotypic traits each other. Node shapes indicate the omics data type belongs to. Red and blue lines indicate positive and negative correlation, respectively. (B) A summary figure illustrating potential mechanisms of *A. muciniphila* to counteract fructose-induced metabolic dysfunction, which was inferred from present multiomcs study and literature evidence. *A. muciniphila* shapes favorable environment for other beneficiary bacteria to thrive and stimulates bile acids production which leads to OEA biosynthesis in the gut. OEA is proposed as a mediator between gut-brain axis, which induces pathways including AVP and OXT signaling, thermogenesis, and insulin regulation to regulate energy homeostasis.

Between microbiota and metabolome, we observed significant correlations between different bacterial families and bile acids, which supports the important role of microbiome in regulating the bile acid pool ^71^. Primary bile acids that are produced in the liver can be further modified into secondary bile acids in the gut, and both steps are regulated by bacteria ^72^. In our network, four primary bile acids (chenodeoxycholate, taurochenodeoxycholate, chenodeoxycholic acid sulfate, cholate sulfate) and three secondary bile acids (hyocholate, taurochenodeoxycholate, 6-beta-hydroxylithicholate) were directly correlated with *Akkermansiaceae*, suggesting a role of *Akkermansiaceae* in bile acid regulation. Hagi et al. reported that the ratio of primary and secondary bile acids is important for the growth of *A. muciniphila* in the culture medium ^73^. This suggests there is a potential environmental reconstruction feedback loop, where *A. muciniphila* creates more suitable growth conditions for itself by recruiting other bile acid metabolizing bacteria such as *Lactobacillus*, *Clostridium* and *Enterococcus* ^73,74^. In our network, the metabolite OEA, whose biosynthesis is modulated by bile acids ^75,76^, was also positively correlated with *Akkermansiaceae*. Lastly, *Oxt* expression in the hypothalamus was positively but indirectly correlated with *Akkermansiaceae* via inhibition of pathogenic *Peptostreptococcaceae* and *Aerococcaceae*.

Taken together, these integrative multiomic results suggest a molecular cascade linking intestinal *A. muciniphila* to altered hypothalamic signaling (Figure 5B). *A. muciniphila* orchestrates with other beneficial bacteria (e.g., *Lactobacillus*, *Clostridium* and *Enterococcus*) to affect the bile acid pool, increase OEA substrate (phosphatidylethanolamine and phosphatidylcholine) metabolism, and induce OEA biosynthesis in the intestine through the upregulation of NAPE-PLD. We note that *A. muciniphila* activated PPARα and GPR119, known receptors of OEA in the intestine, which can transmit signals to the hypothalamus via the vagus nerve ^77, 78,79^. We also observed that *A. muciniphila* activated hypothalamic oxytocin and vasopressin signaling as well as energy metabolism pathways (thermogenesis and oxidative phosphorylation), all of which can regulate energy and metabolic homeostasis in fructose-induced MetS. Based on these results, we hypothesize that OEA may mediate the effects of *A. muciniphila*.

### Administration of OEA partially recapitulates the metabolic benefits of *A. mucinipihla*

To validate whether OEA is one of the metabolites mediating the beneficial effects of *A. muciniphila*, we orally gavaged OEA (or vehicle) into DBA mice with or without fructose feeding and examined phenotypes and gene expressions in small intestine and hypothalamus (Figure 6A). We observed a significant decrease of food intake and increase of liquid intake in fructose fed mice; however, OEA did not modify these effects. Overall calorie intakes were the same across all groups (Figure 6B,C). Decreased food intake by OEA has been observed in previous studies in control diet-fed rats and high-fat diet mice ^80,81^, but in our fructose diet study OEA did not influence food intake, demonstrating that the effect of OEA on food intake is dependent on dietary condition. OEA treatment significantly mitigated fructose-induced body weight gain over 8 weeks as well as glucose intolerance (p = 0.036 and p = 0.021 by Two-way repeated measures ANOVA test, respectively; Figure 6D,E). OEA also non-significantly dampened the fructose-induced increases in total and individual fat mass including gonadal and inguinal white adipose tissues (gWAT and iWAT) (Figure 6F). To test whether thermogenesis was involved in the effects of OEA on body weight, we measured body temperature at week 7. OEA significantly increased rectal temperature compared to vehicle under fructose consumption, supporting that OEA induces thermogenesis and energy expenditure (Figure 6G).

**Figure 6.**
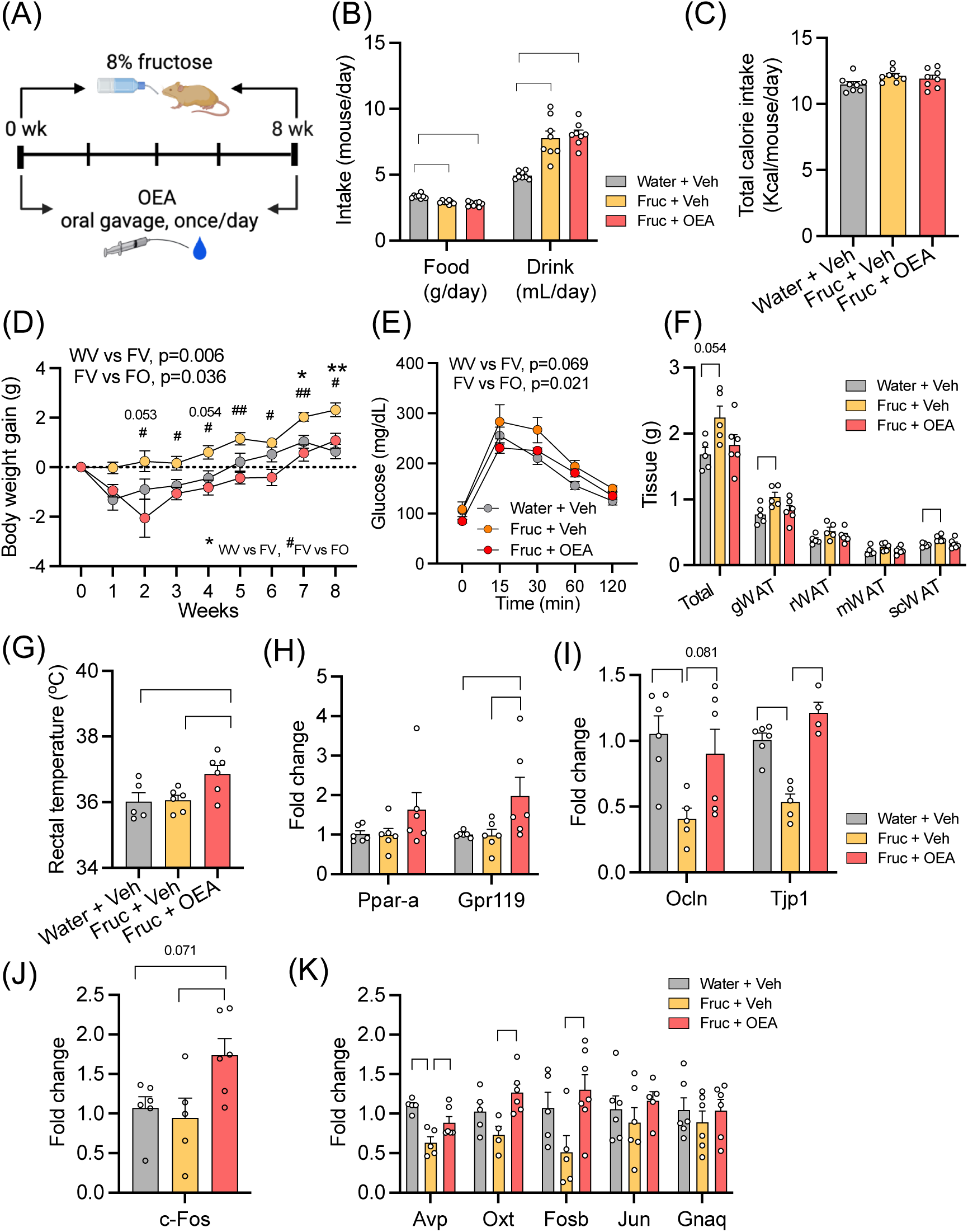
OEA administration partially recapitulates metabolic benefit of *A. muchiniphila.* (A) Overall experimental design of OEA administration. (B-C) Food, drink, and calorie intake of DBA mice during 8 weeks of OEA administration with fructose or normal water. (D) Body weight gain over 8 weeks. (E) glucose levels during intraperitoneal glucose-tolerance test (IPGTT) at week 8. (D-E) On the top of the graph, the p-value of main effects (fructose, OEA) are shown for the comparisons of Water + Veh vs. Fructose + Veh (WV vs. FV, fructose effect) or Fructose + Veh vs. Fructose + OEA (FV vs. FO, OEA effect). Asterisks (*) or number (#) signs indicate time points at which significant differences were found in WV vs. FV or FV vs. FO, respectively. Statistical significance was determined by Two-way repeated measures ANOVA with Tukey’s post hoc test. ^#,^ * p-val < 0.05, ** p-val < 0.01. (F) Adipose depots, gWAT, rWAT, mWAT, scWAT: gonadal, retroperitoneal, mesenteric, subcutaneous white adipose tissue, respectively. Total was calculated by summing these four individual adipose depots. (G) Rectal temperature measurement at week 7. (H) Fold changes of expression level of OEA receptor genes (*Pparα* and *Gpr119*). (I) Fold changes of expression level of tight junction related gene, *Ocln* and *Tjp1* in the ileum. (J-K) Fold changes of expression level of hypothalamic genes (*c-Fos, Avp, Oxt, Fosb, Jun*, and *Gnaq)*. Data are presented as means ± SEMs. n=5-8/group. (B-C), (F-K), Statistical significance was determined by One-way ANOVA with Tukey’s post hoc test. P-values were considered statistically significant at p-val < 0.05. * p-val < 0.05, ** p-val < 0.01, *** p-val < 0.001

To determine whether OEA administration recapitulates the effects of *A. muciniphila* on intestinal and hypothalamic gene expression, we performed real-time qPCR analysis. In ileum, OEA treatment induced a significant increase in *Gpr119* expression and a non-significant increasing trend in *Pparα* expression compared with vehicle under fructose feeding (Figure 6H). We also tested the tight junction-related genes, *Ocln* encoding occludin and *Tjp1* encoding tight junction protein 1 (ZO-1). Expression of both genes were significantly decreased by fructose diet and OEA reversed the fructose effect (Figure 6I). In hypothalamus, we also tested genes involved in oxytocin and vasopressin signaling which were found to be DEGs in our hypothalamus scRNA-seq analysis for *A. muciniphila* (*Oxt*, *Jun*, c-*Fos*, *Fosb*, *Jun*, *Gnaq, Avp*). Among these, OEA significantly increased hypothalamic expression of *c-Fos*, a key immediate early gene and a marker for neural activity ^53^, *Avp*, and genes involved in oxytocin signaling including *Oxt* and *Fosb* (Figure 6J,K).

Taken together, the metabolic phenotypes and gene expression patterns altered by OEA partially recapitulate the effects of *A. muciniphila*, with consistencies in decreasing body weight and glucose intolerance, increasing intestinal OEA receptors and tight junction markers, and activating hypothalamic genes involved in neuronal activation and vasopressin and oxytocin signaling.

### Comparison of convergent and divergent effects of *A. muciniphila* between fructose and high-fat diet mouse models

To elucidate whether the effects and mechanisms of *A. muciniphila* observed in our fructose-induced MetS model converge or diverge from those in the widely used HFD-induced MetS model, we compared our results with previously published data on the HFD mouse model at phenotypic, gut microbiome, metabolomics, and transcriptome levels (Table 1).

**Table 1.**
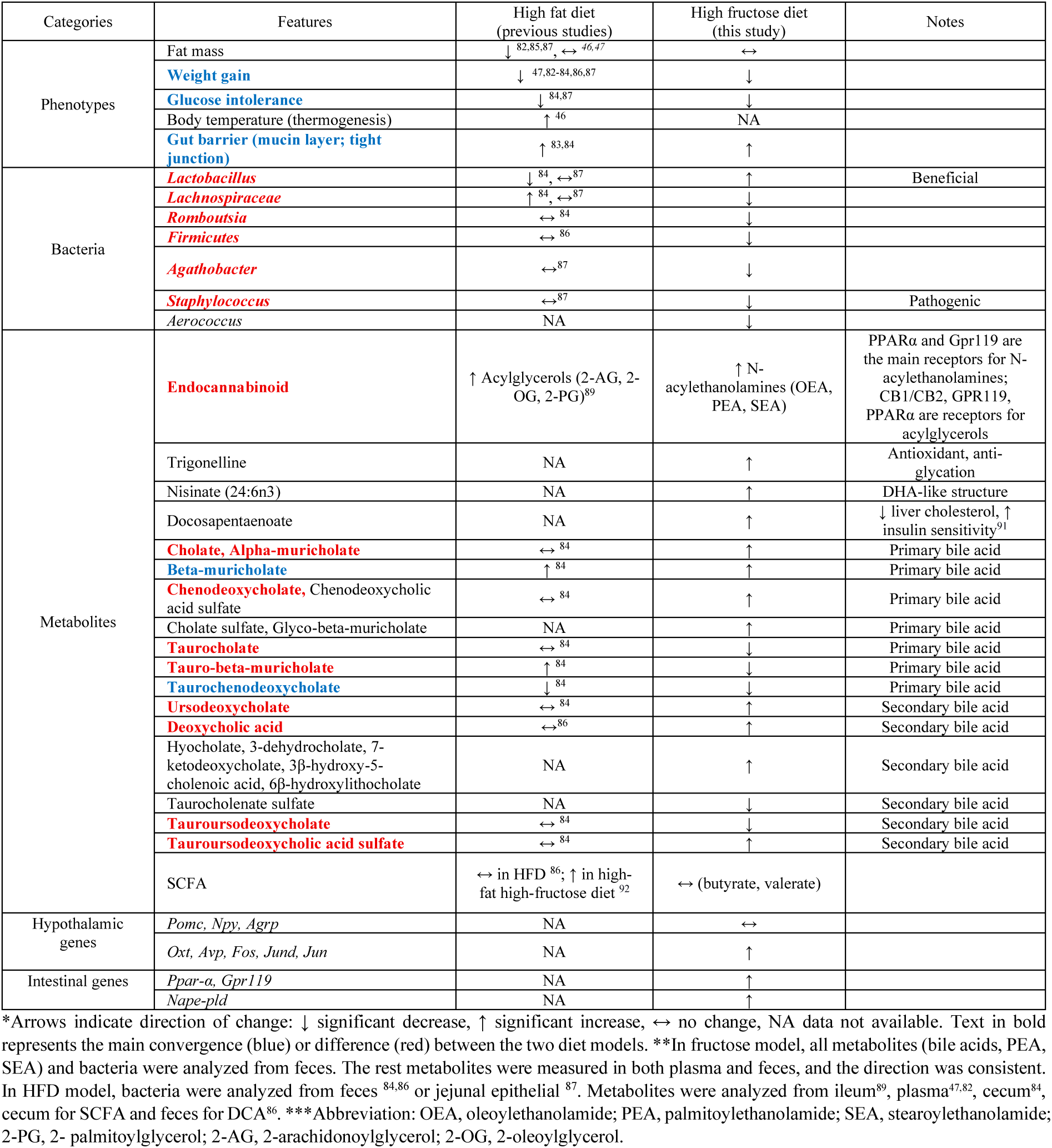
Comparison of features differentially regulated by *A. muciniphila* treatment under high-fat or fructose diet.

At the phenotypic level, *A. muciniphila* improved metabolic phenotypes in both models by decreasing weight gain, increasing glucose sensitivity, and improving gut barrier function in both fructose and HFD MetS models ^47,82–87^, but the effect on fat mass was less consistent.

At the microbiome level, in our fructose MetS model *A. muciniphila* mainly increased *Lactobacillus* and decreased *Lachnospiraceae*, but these taxa exhibited opposite trends or no change in the HFD model. *A. muciniphila* also uniquely decreased *Romboutsia, Agathobactor, Fermicutes,* and *Staphylococcus* in the fructose model.

At the metabolomics level, *A. muciniphila* increased beta-muricholate and decreased taurochenodeoxycholate in both HFD and fructose diets, but several other major primary bile acids (chenodeoxycholate and alpha-muricholate) and secondary bile acids (e.g., ursodeoxycholate) were increased by *A. muciniphila* only in fructose MetS. All other tau-conjugated bile acids such as tauro-beta-muricholate and tauroursodeoxycholate were decreased by *A. muciniphila* in fructose diet, but were either increased or unchanged under HFD ^84^. Apart from bile acids, *A. muciniphila* increased endocannabinoids in feces under both diets, but the class of endocannabinoids varied according to the diet: acylglycerols (2-PG, 2-AG, 2-OG; activate cannabinoid receptors (CB1/CB2) and exhibit more variable, and often pro-lipogenic, effects on metabolic regulation^88^) in HFD ^89^ versus endocannabinoid-like N-acylethanolamines (OEA, PEA, SEA; ligand of PPAR-a and associated with regulation of lipid metabolism and energy homeostasis) in high fructose. Trigonelline, a methylated form of vitamin B3 previously reported to inhibit hepatic lipid accumulation^90^, as well as long-chain omega-3 fatty acids involved in DHA biosynthesis, docosapentaenoate (DPA) and nisinate, were significantly increased by *A. muciniphila* in fructose model; however, its effects on these metabolites have not been reported in HFD model despite evidence that DPA exerts beneficial effects ^91^. SCFA production by *A. muciniphila*, especially acetate and propionate, is well documented in vitro and is hypothesized to mediate part of its metabolic benefits. In a high-fat high-fructose diet (HFHFD) model, *A. muciniphila* increased SCFAs in the stool and liver ^92,93,94^. However, in both HFD ^86^ and our fructose models, SCFA was not affected by *A. muciniphila*.

Overall, our systematic comparison of the *A. muciniphila* effects between our fructose MetS model and the HFD model reveals convergence on improving body weight, glucose sensitivity, and intestinal tight junctions, but highlights remarkable differences in gut bacteria and metabolites between the two models. For hypothalamic or intestinal genes, direct comparison between the two dietary models was not feasible due to the limited availability of reports of *A. muciniphila* effect on the brain and intestine in the HFD model.

## Discussion

In the present study, using a multitissue multiomics systems biology approach coupled with experimental validation, we unveiled the molecular cascades involved in the beneficial effect of *A. muciniphila* on fructose-induced MetS. At the microbiota level, *A. muciniphila* treatment promoted the proliferation of beneficial bacteria such as *Lactobacillus, Bacteroides*,

*Clostridium*, and *Enterococcus* while inhibiting the growth of pathogenic taxa including *Staphylococcus* and *Aerococcus*. Global metabolomics revealed major increases in bile acids (both primary and secondary) and the endocannabinoid analogue OEA, accompanied by reductions in tau-conjugated bile acids, consistent with the activity of *Lactobacillus*, *Bacteroides*, *Clostridium*, and *Enterococcus* species known to deconjugate bile acids ^67,72,76,95–98^. scRNA-seq further revealed hypothalamic alterations in pathways linked to energy metabolism and endocannabinoid signaling across neuronal and glial populations. Experimental validation confirmed that OEA partially mimics *A. muciniphila* effects in improving fructose-induced obesity and glucose intolerance, reinforcing gut barrier function and activating intestinal receptor signaling, and stimulating oxytocin and vasopressin signaling in the hypothalamus, thereby supporting OEA as a functional regulator of *A. muciniphila*’s beneficial effects in the gut-brain axis. Together, these data support a gut–brain cascade from *A. muciniphila* to bile acids to OEA to hypothalamic reprogramming in counteracting fructose-induced MetS.

Previous work has highlighted diverse mechanisms underlying the metabolic benefits of *A. muciniphila*. For instance, under high-fat feeding, Chelakkot et al. showed that *A. muciniphila* strengthens gut barrier integrity through TLR2 signaling and extracellular vesicles ^99^, thereby reducing lipopolysaccharide leakage and pro-inflammatory cytokines such as IFNγ and TNF-α ^100,101^. It also converts dietary fiber into SCFAs that regulate glucose and lipid metabolism and has been linked to serotonin elevation in the colon and serum ^49,93^.

In our fructose model, we observed parallel improvements in gut barrier function and dampened hypothalamic inflammation by *A. muciniphila* (Figure 1I,J, Figure S8). Reduced caloric absorption through enhanced gut barrier function and transit ^102^ may partially explain the reduced overall body weight. Additionally, activation of hypothalamic energy-expenditure pathways (thermogenesis, oxidative phosphorylation) suggests that elevated energy expenditure drives weight reduction ^89,103^. Our scRNA-seq analysis indeed supports that *A. muciniphila* reprograms hypothalamic signaling through both neuronal and non-neuronal cells. Oxytocin (OXT) and vasopressin (AVP) pathways were among the most strongly altered, with cell-type and diet specificity. Under fructose feeding, OXT signaling and AVP-regulated water reabsorption were enriched in neurons as well as other cell types. These neuropeptides are known to increase energy expenditure, lipolysis, insulin sensitivity, and thermogenesis, supporting the observed improvements in metabolic outcomes ^52,104,105^. Gene-level changes reinforced these pathway alterations: immediate-early genes (*Fos*, *Jun*) and neuromodulators (*Gal*, *Meg3*) were also induced, consistent with OXT/AVP-driven neuronal activation and glucose homeostasis regulation. Among the neuronal subtypes, *Fos* upregulation was the strongest in the Pdyn/Oxt and Npy/Agrp subtypes particularly in the fructose group, suggesting *A. muciniphila* may activate neurons related to energy homeostasis. Other DEGs such as *Pkhd1*, *Ndfip1*, and *Gas5* are associated with ciliary signaling, proteostasis, and stress adaptation, respectively. Together, these results demonstrate that *A. muciniphila* exerts central metabolic benefits through diet-specific modulation of hypothalamic neuropeptides and diverse signaling pathways central to energy homeostasis.

In searching for the regulators of *A. muciniphila-*to-hypothalamus crosstalk, a central discovery from our plasma and fecal metabolomics analysis is the coordinated regulation of bile acid metabolism and OEA synthesis by *A. muciniphila* in fructose-induced MetS. We observed significant increases in both primary (CDCA, UDCA) and secondary bile acids. In mice, CDCA and UDCA are usually at low levels due to the rapid conversion into muricholic acids, but both were strongly elevated together with muricholic acids under *A. muciniphila* treatment, suggesting enhanced hepatic synthesis and/or incomplete conversion ^106^.

Mechanistically, CDCA has the highest affinity for NAPE-PLD, the key OEA-synthesizing enzyme in gut epithelial cells ^67^. Consistent with this, *A. muciniphila* upregulated expression of NAPE-PLD, an enzyme involved in OEA biosynthesis, and PPARα and GPR119, the receptors through which OEA acts to activate gut–brain signaling. While OEA shares structural similarity with endocannabinoids, it does not activate CB1/CB2 ^107^, instead triggering PPARα- and GPR119-mediated signaling to promote oxytocin release and hypothalamic signaling reprogramming ^108,109^. Additional metabolite changes by *A. muciniphila* under fructose diet, including increased vitamin B5 and vitamin K, further point to potential complementary mechanisms which activate brown or white adipose tissue, respectively, and modulate glucose/fat metabolism ^110,111^.

Connecting the multiomics data together, our network analysis provides a systems-level perspective of *A. muciniphila*’s actions. The bacterium fosters a favorable microbial environment, enriching *Lactobacillaceae*, *Erysipelotrichaceae*, and *Clostridiaceae* while inhibiting pathogenic taxa ^112–114^. Most bile acids were positively correlated with *A. muciniphila* treatment, recapitulating their mutual reinforcement ^106^. Importantly, OEA emerged as the most coherent metabolite linking gut microbiota changes to hypothalamic gene regulation via PPARα and GPR119 ^75,76,115–117^. This places OEA at the core of the *A. muciniphila*–driven gut–brain axis.

While decreasing weight gain, increasing glucose sensitivity, and gut barrier improvement are a common effect of *A. muciniphila* in both fructose and HFD models, the microbiota and metabolite responses significantly differ (Table 1). In the fructose model, *A. muciniphila* increased *Lactobacillus* and decreased *Staphylococcus*, *Aerococcus*, and *Romboutsia*, whereas in HFD feeding, *Lactobacillus* was primarily reduced ^84,118^. Metabolite responses also diverged: although some bile acids (e.g., taurochenodeoxycholate, beta-muricholate) shifted similarly, others (e.g., tauro-beta-muricholate, tauroursodeoxycholic acid sulfate) moved in opposite directions. Most notably, *A. muciniphila* regulated NAEs endocannabinoid analogues (OEA, PEA, SEA) in the fructose model, whereas acylglycerols dominated in the HFD study ^14^. This distinction is significant because acylglycerols are primarily ligands for the CB1 receptor, which has been associated with psychiatric adverse effects. In contrast, endocannabinoid analogues such as OEA do not act through the CB1 receptor and have emerged as potentially safer anti-obesity therapeutics ^51,107^. These results highlight MetS subtype- specific molecular mechanisms and identify OEA as a key mediator of metabolic reprogramming by *A. muciniphila* under fructose-induced MetS. Taken together, these results point to the versatile functions of *A. muciniphila* to modulate different metabolites with therapeutic effects in different disease settings and offer precision medicine targets for different MetS subtypes.

We acknowledge the following limitations. Causal inference across microbes, metabolites, and host genes requires deeper experimental validation. Although our OEA experiments confirmed activation of gut and brain signaling pathways, whether OEA acts mainly through the vagus nerve or also directly via brain entry remains to be clarified. Caution for translational generalization to humans is also needed. In a human MetS cohort without specific dietary intervention, *A. muciniphila* did not significantly increase overall circulating endocannabinoids but selectively increased 1-PG and 2-PG ^119^, which differs from the OEA increase we found under fructose-induced MetS and the previously reported increases in acylglycerols (2-AG, 2-OG, 2-PG) in HFD induced MetS in mice ^14^. Broader testing across diets, hosts, and disease conditions will be critical to identify universal versus condition-specific mediators.

In summary, our multiomics study delineates how *A. muciniphila* confers metabolic benefits in fructose-induced MetS in mice. By shaping the gut microbiota, remodeling bile acid pools, and enhancing OEA synthesis, *A. muciniphila* triggers gut–brain interactions that reprogram hypothalamic energy metabolism and neuropeptide signaling. These findings not only provide mechanistic understanding but also point to the endocannabinoid OEA and bile acid pathways as diet-specific therapeutic targets for fructose-associated metabolic disorders.

## Acknowledgments

This work was supported by NIH DK104363 (XY) and QCB Collaboratory Fellowship (S.M.H.). We thank Wini Suryavanshi for her valuable assistance with the animal experiments.

## Author contributions

X.Y., S.M.H. and I.S.A. conceived the study design. S.M.H. and I.S.A. drafted the manuscript and generated figures. S.M.H., X.Y., E.YH., I.S.A., and S.W. reviewed and edited manuscript, which was approved by all authors. I.S.A., J.Y., T.K., CAO, Z.Z., R.L. and A.C. conducted animal experiment and qPCR experiment. I.S.A., G.D., G.Z., and I.C. performed Drop-seq and library preparation for scRNA-seq. I.S.A and J.Y. performed metabolomics data analyses. S.M.H. performed the bioinformatic analyses of scRNA-seq, microbiome data, and multiomics data integration.

## Ethics approval statement

The ethics application (Approval No: ARC-2012-059) for mouse study was approved by the UCLA Institutional Animal Care and Use Committee.

## Lead Contact and Materials Availability

Further information and requests for resources and code should be directed to and will be fulfilled by the Lead Contact, Xia Yang **.**

## Data availability statement

scRNA-seq data have been deposited at NCBI Gene Expression Omnibus (GEO) database (Accession: GSE212491). Raw 16S rRNA sequencing data for all samples have been deposited in the open-source repository NCBI Sequence Read Archive (SRA) database (Accession: PRJNA876737). All mass spectrometry data analyzed in this study have been deposited to Mendeley Data and are publicly available (URL: https://doi.org/10.17632/f55rsncm9y.1). All the reagents and primers used in this study are provided in Table S5.

## References

1. Goncalves, M.D., Lu, C., Tutnauer, J., Hartman, T.E., Hwang, S.-K., Murphy, C.J., Pauli, C., Morris, R., Taylor, S., Bosch, K., et al. (2019). High-fructose corn syrup enhances intestinal tumor growth in mice. Science (New York, N.Y.) 363, 1345–1349. 10.1126/science.aat8515.

2. Johnson, R.J., Perez-Pozo, S.E., Sautin, Y.Y., Manitius, J., Sanchez-Lozada, L.G., Feig, D.I., Shafiu, M., Segal, M., Glassock, R.J., Shimada, M., et al. (2009). Hypothesis: could excessive fructose intake and uric acid cause type 2 diabetes? Endocr Rev 30, 96–116. 10.1210/er.2008-0033.

3. Sindhunata, D.P., Meijnikman, A.S., Gerdes, V.E.A., and Nieuwdorp, M. (2022). Dietary fructose as a metabolic risk factor. American Journal of Physiology-Cell Physiology. 10.1152/ajpcell.00439.2021.

4. Stanhope, K.L., and Havel, P.J. (2008). Endocrine and metabolic effects of consuming beverages sweetened with fructose, glucose, sucrose, or high-fructose corn syrup. Am J Clin Nutr 88, 1733S–1737S. 10.3945/ajcn.2008.25825d.

5. Grundy, S.M., Cleeman, J.I., Daniels, S.R., Donato, K.A., Eckel, R.H., Franklin, B.A., Gordon, D.J., Krauss, R.M., Savage, P.J., Smith, S.C., Jr., et al. (2005). Diagnosis and management of the metabolic syndrome: an American Heart Association/National Heart, Lung, and Blood Institute Scientific Statement. Circulation 112, 2735–2752. 10.1161/CIRCULATIONAHA.105.169404.

6. Kim, M.S., Krawczyk, S.A., Doridot, L., Fowler, A.J., Wang, J.X., Trauger, S.A., Noh, H.L., Kang, H.J., Meissen, J.K., Blatnik, M., et al. (2016). ChREBP regulates fructose-induced glucose production independently of insulin signaling. Journal of Clinical Investigation 126, 4372–4386. 10.1172/JCI81993.

7. Meng, Q., Ying, Z., Noble, E., Zhao, Y., Agrawal, R., Mikhail, A., Zhuang, Y., Tyagi, E., Zhang, Q., Lee, J.-H., et al. (2016). Systems Nutrigenomics Reveals Brain Gene Networks Linking Metabolic and Brain Disorders. 10.1016/j.ebiom.2016.04.008.

8. Zhang, G., Byun, H.R., Ying, Z., Blencowe, M., Zhao, Y., Hong, J., Shu, L., Chella Krishnan, K., Gomez-Pinilla, F., and Yang, X. (2020). Differential metabolic and multi-tissue transcriptomic responses to fructose consumption among genetically diverse mice. Biochimica et Biophysica Acta - Molecular Basis of Disease. 10.1016/j.bbadis.2019.165569.

9. Chen, Y.W., Ahn, I.S., Wang, S.S., Majid, S., Diamante, G., Cely, I., Zhang, G., Cabanayan, A., Komzyuk, S., Bonnett, J., et al. (2025). Multitissue single-cell analysis reveals differential cellular and molecular sensitivity between fructose and high-fat high-sucrose diets. Cell Rep 44, 115690. 10.1016/j.celrep.2025.115690.

10. Do, M., Lee, E., Oh, M.-J., Kim, Y., and Park, H.-Y. (2018). High-Glucose or -Fructose Diet Cause Changes of the Gut Microbiota and Metabolic Disorders in Mice without Body Weight Change. Nutrients 10, 761–761. 10.3390/nu10060761.

11. Jang, C., Hui, S., Lu, W., Tesz, G.J., Birnbaum, M.J., Rabinowitz, J.D., Jang, C., Hui, S., Lu, W., Cowan, A.J., et al. (2018). The Small Intestine Converts Dietary Fructose into Glucose and Organic Acids. Cell Metabolism 27, 351–361.e353. 10.1016/j.cmet.2017.12.016.

12. Ahn, I.S., Lang, J.M., Olson, C.A., Diamante, G., Zhang, G., Ying, Z., Byun, H.R., Cely, I., Ding, J., Cohn, P., et al. (2020). Host genetic background and gut microbiota contribute to differential metabolic responses to fructose consumption in mice. Journal of Nutrition. 10.1093/jn/nxaa239.

13. Ramirez, C.B., Ahn, I.S., Rubtsova, V.I., Cely, I., Le, J., Kim, J., Jung, S., Kelly, M.E., Kim, Y., Bae, H., et al. (2025). Circulating glycerate predicts resilience to fructose-induced hepatic steatosis. Cell Metab 37, 1223–1234 e1225. 10.1016/j.cmet.2025.03.017.

14. Reikvam, D.H., Erofeev, A., Sandvik, A., Grcic, V., Jahnsen, F.L., Gaustad, P., McCoy, K.D., Macpherson, A.J., Meza-Zepeda, L.A., and Johansen, F.-E. (2011). Depletion of murine intestinal microbiota: effects on gut mucosa and epithelial gene expression. PloS one 6, e17996–e17996. 10.1371/journal.pone.0017996.

15. Olson, C.A., Vuong, H.E., Yano, J.M., Liang, Q.Y., Nusbaum, D.J., and Hsiao, E.Y. (2018). The Gut Microbiota Mediates the Anti-Seizure Effects of the Ketogenic Diet. Cell 173, 1728–1741.e1713. 10.1016/j.cell.2018.04.027.

16. Zhang, Z., Park, J.W., Ahn, I.S., Diamante, G., Sivakumar, N., Arneson, D., Yang, X., van Veen, J.E., and Correa, S.M. (2021). Estrogen receptor alpha in the brain mediates tamoxifen-induced changes in physiology in mice. Elife 10. 10.7554/eLife.63333.

17. Bridgewater Br, E.A.M. (2014). High Resolution Mass Spectrometry Improves Data Quantity and Quality as Compared to Unit Mass Resolution Mass Spectrometry in High-Throughput Profiling Metabolomics. Journal of Postgenomics Drug & Biomarker Development 04. 10.4172/2153-0769.1000132.

18. Ford, L., Kennedy, A.D., Goodman, K.D., Pappan, K.L., Evans, A.M., Miller, L.A.D., Wulff, J.E., Wiggs Iii, B.R., Lennon, J.J., Elsea, S., and Toal, D.R. (2020). Precision of a Clinical Metabolomics Profiling Platform for Use in the Identification of Inborn Errors of Metabolism. The Journal of Applied Laboratory Medicine 5, 342–356. 10.1093/jalm/jfz026.

19. Zierer, J., Jackson, M.A., Kastenmüller, G., Mangino, M., Long, T., Telenti, A., Mohney, R.P., Small, K.S., Bell, J.T., Steves, C.J., et al. (2018). The fecal metabolome as a functional readout of the gut microbiome. Nature Genetics 50, 790–795. 10.1038/s41588-018-0135-7.

20. Brewer, G.J., and Torricelli, J.R. (2007). Isolation and culture of adult neurons and neurospheres. Nature protocols 2, 1490–1498. 10.1038/nprot.2007.207.

21. Arneson, D., Zhang, G., Ying, Z., Zhuang, Y., Byun, H.R., Ahn, I.S., Gomez-Pinilla, F., and Yang, X. (2018). Single cell molecular alterations reveal target cells and pathways of concussive brain injury. Nature Communications 9, 3894–3894. 10.1038/s41467-018-06222-0.

22. Macosko, E.Z., Basu, A., Satija, R., Nemesh, J., Shekhar, K., Goldman, M., Tirosh, I., Bialas, A.R., Kamitaki, N., Martersteck, E.M., et al. (2015). Highly Parallel Genome-wide Expression Profiling of Individual Cells Using Nanoliter Droplets. Cell 161, 1202–1214. 10.1016/j.cell.2015.05.002.

23. Wang, X., Wang, X., and Wang, T. (2012). Synthesis of oleoylethanolamide using lipase. J Agric Food Chem 60, 451–457. 10.1021/jf203629w.

24. Reagan-Shaw, S., Nihal, M., and Ahmad, N. (2008). Dose translation from animal to human studies revisited. FASEB journal : official publication of the Federation of American Societies for Experimental Biology 22, 659–661. 10.1096/fj.07-9574LSF.

25. Spandidos, A., Wang, X., Wang, H., and Seed, B. (2010). PrimerBank: a resource of human and mouse PCR primer pairs for gene expression detection and quantification. Nucleic Acids Research 38, D792–D799. 10.1093/nar/gkp1005.

26. Bolyen, E., Rideout, J.R., Dillon, M.R., Bokulich, N.A., Abnet, C.C., Al-Ghalith, G.A., Alexander, H., Alm, E.J., Arumugam, M., Asnicar, F., et al. (2019). Reproducible, interactive, scalable and extensible microbiome data science using QIIME 2. Nature Biotechnology 37, 852–857. 10.1038/s41587-019-0209-9.

27. Callahan, B.J., McMurdie, P.J., Rosen, M.J., Han, A.W., Johnson, A.J.A., and Holmes, S.P. (2016). DADA2: High-resolution sample inference from Illumina amplicon data. Nature Methods 13, 581–583. 10.1038/nmeth.3869.

28. Quast, C., Pruesse, E., Yilmaz, P., Gerken, J., Schweer, T., Yarza, P., Peplies, J., and Glöckner, F.O. (2013). The SILVA ribosomal RNA gene database project: improved data processing and web-based tools. Nucleic Acids Research 41, D590–D596. 10.1093/nar/gks1219.

29. Segata, N., Izard, J., Waldron, L., Gevers, D., Miropolsky, L., Garrett, W.S., and Huttenhower, C. (2011). Metagenomic biomarker discovery and explanation. Genome Biology 12, R60–R60. 10.1186/gb-2011-12-6-r60.

30. Douglas, G.M., Maffei, V.J., Zaneveld, J.R., Yurgel, S.N., Brown, J.R., Taylor, C.M., Huttenhower, C., and Langille, M.G.I. (2020). PICRUSt2 for prediction of metagenome functions. Nature Biotechnology 38, 685–688. 10.1038/s41587-020-0548-6.

31. DeHaven, C.D. (2012). Software Techniques for Enabling High-Throughput Analysis of Metabolomic Datasets. In A.M. Evans, ed. (IntechOpen), pp. Ch. 7–Ch. 7. 10.5772/31277.

32. Ahn, I.S., Yoon, J., Diamante, G., Cohn, P., Jang, C., and Yang, X. (2021). Disparate Metabolomic Responses to Fructose Consumption between Different Mouse Strains and the Role of Gut Microbiota. Metabolites 11. 10.3390/metabo11060342.

33. DePasquale, E.A.K., Schnell, D.J., Van Camp, P.-J., Valiente-Alandí, Í., Blaxall, B.C., Grimes, H.L., Singh, H., and Salomonis, N. (2019). DoubletDecon: Deconvoluting Doublets from Single-Cell RNA-Sequencing Data. Cell Reports 29, 1718–1727.e1718. 10.1016/j.celrep.2019.09.082.

34. Romanov, R.A., Zeisel, A., Bakker, J., Girach, F., Hellysaz, A., Tomer, R., Alpár, A., Mulder, J., Clotman, F., Keimpema, E., et al. (2017). Molecular interrogation of hypothalamic organization reveals distinct dopamine neuronal subtypes. Nature Neuroscience 20, 176–188. 10.1038/nn.4462.

35. Chen, R., Wu, X., Jiang, L., and Zhang, Y. (2017). Single-Cell RNA-Seq Reveals Hypothalamic Cell Diversity. Cell reports 18, 3227–3241. 10.1016/j.celrep.2017.03.004.

36. Hammond, T.R., Dufort, C., Dissing-Olesen, L., Giera, S., Young, A., Wysoker, A., Walker, A.J., Gergits, F., Segel, M., Nemesh, J., et al. (2019). Single-Cell RNA Sequencing of Microglia throughout the Mouse Lifespan and in the Injured Brain Reveals Complex Cell-State Changes. Immunity 50, 253–271.e256. 10.1016/j.immuni.2018.11.004.

37. Choudhary, S., and Satija, R. (2022). Comparison and evaluation of statistical error models for scRNA-seq. Genome Biology 23, 27–27. 10.1186/s13059-021-02584-9.

38. Xie, Z., Bailey, A., Kuleshov, M.V., Clarke, D.J.B., Evangelista, J.E., Jenkins, S.L., Lachmann, A., Wojciechowicz, M.L., Kropiwnicki, E., Jagodnik, K.M., et al. (2021). Gene Set Knowledge Discovery with Enrichr. Current Protocols 1, e90–e90. 10.1002/cpz1.90.

39. Friedman, J., and Alm, E.J. (2012). Inferring correlation networks from genomic survey data. PLoS computational biology 8, e1002687–e1002687. 10.1371/journal.pcbi.1002687.

40. Watts, S.C., Ritchie, S.C., Inouye, M., and Holt, K.E. (2019). FastSpar: rapid and scalable correlation estimation for compositional data. Bioinformatics (Oxford, England) 35, 1064–1066. 10.1093/bioinformatics/bty734.

41. Langfelder, P., and Horvath, S. (2008). WGCNA: an R package for weighted correlation network analysis. BMC Bioinformatics 9, 559–559. 10.1186/1471-2105-9-559.

42. Arneson, D., Bhattacharya, A., Shu, L., Mäkinen, V.-P., and Yang, X. (2016). Mergeomics: a web server for identifying pathological pathways, networks, and key regulators via multidimensional data integration. BMC genomics 17, 722–722. 10.1186/s12864-016-3057-8.

43. Ding, J., Blencowe, M., Nghiem, T., Ha, S.-M., Chen, Y.-W., Li, G., and Yang, X. (2021). Mergeomics 2.0: a web server for multi-omics data integration to elucidate disease networks and predict therapeutics. Nucleic acids research. 10.1093/nar/gkab405.

44. Shu, L., Zhao, Y., Kurt, Z., Byars, S.G., Tukiainen, T., Kettunen, J., Orozco, L.D., Pellegrini, M., Lusis, A.J., Ripatti, S., et al. (2016). Mergeomics: multidimensional data integration to identify pathogenic perturbations to biological systems. BMC Genomics 17, 874–874. 10.1186/s12864-016-3198-9.

45. Shannon, P., Markiel, A., Ozier, O., Baliga, N.S., Wang, J.T., Ramage, D., Amin, N., Schwikowski, B., and Ideker, T. (2003). Cytoscape: a software environment for integrated models of biomolecular interaction networks. Genome research 13, 2498–2504. 10.1101/gr.1239303.

46. Yoon, H.S., Cho, C.H., Yun, M.S., Jang, S.J., You, H.J., Kim, J.H., Han, D., Cha, K.H., Moon, S.H., Lee, K., et al. (2021). Akkermansia muciniphila secretes a glucagon-like peptide-1-inducing protein that improves glucose homeostasis and ameliorates metabolic disease in mice. Nat Microbiol 6, 563–573. 10.1038/s41564-021-00880-5.

47. Plovier, H., Everard, A., Druart, C., Depommier, C., Van Hul, M., Geurts, L., Chilloux, J., Ottman, N., Duparc, T., Lichtenstein, L., et al. (2017). A purified membrane protein from Akkermansia muciniphila or the pasteurized bacterium improves metabolism in obese and diabetic mice. Nat Med 23, 107–113. 10.1038/nm.4236.

48. Xu, R., Zhang, Y., Chen, S., Zeng, Y., Fu, X., Chen, T., Luo, S., and Zhang, X. (2023). The role of the probiotic Akkermansia muciniphila in brain functions: insights underpinning therapeutic potential. Crit Rev Microbiol 49, 151–176. 10.1080/1040841x.2022.2044286.

49. Yaghoubfar, R., Behrouzi, A., Ashrafian, F., Shahryari, A., Moradi, H.R., Choopani, S., Hadifar, S., Vaziri, F., Nojoumi, S.A., Fateh, A., et al. (2020). Modulation of serotonin signaling/metabolism by Akkermansia muciniphila and its extracellular vesicles through the gut-brain axis in mice. Sci Rep 10, 22119. 10.1038/s41598-020-79171-8.

50. Bennett, M.L., Bennett, F.C., Liddelow, S.A., Ajami, B., Zamanian, J.L., Fernhoff, N.B., Mulinyawe, S.B., Bohlen, C.J., Adil, A., Tucker, A., et al. (2016). New tools for studying microglia in the mouse and human CNS. Proc Natl Acad Sci U S A 113, E1738–1746. 10.1073/pnas.1525528113.

51. Cristino, L., Bisogno, T., and Di Marzo, V. (2020). Cannabinoids and the expanded endocannabinoid system in neurological disorders. Nature reviews. Neurology 16, 9–29. 10.1038/s41582-019-0284-z.

52. Ding, C., and Magkos, F. (2019). Oxytocin and Vasopressin Systems in Obesity and Metabolic Health: Mechanisms and Perspectives. Current obesity reports 8, 301–316. 10.1007/s13679-019-00355-z.

53. Lara Aparicio, S.Y., Laureani Fierro, A.J., Aranda Abreu, G.E., Toledo Cardenas, R., Garcia Hernandez, L.I., Coria Avila, G.A., Rojas Duran, F., Aguilar, M.E.H., Manzo Denes, J., Chi-Castaneda, L.D., and Perez Estudillo, C.A. (2022). Current Opinion on the Use of c-Fos in Neuroscience. NeuroSci 3, 687–702. 10.3390/neurosci3040050.

54. Karatayev, O., Baylan, J., and Leibowitz, S.F. (2009). Increased intake of ethanol and dietary fat in galanin overexpressing mice. Alcohol 43, 571–580. 10.1016/j.alcohol.2009.09.025.

55. Izdebska, K., and Ciosek, J. (2010). Galanin influences on vasopressin and oxytocin release: in vitro studies. Neuropeptides 44, 341–348. 10.1016/j.mce.2020.110903.

56. Cheng, X., Shihabudeen Haider Ali, M.S., Moran, M., Viana, M.P., Schlichte, S.L., Zimmerman, M.C., Khalimonchuk, O., Feinberg, M.W., and Sun, X. (2021). Long non-coding RNA Meg3 deficiency impairs glucose homeostasis and insulin signaling by inducing cellular senescence of hepatic endothelium in obesity. Redox biology 40, 101863–101863. 10.1016/j.redox.2021.101863.

57. Stojanov, S., Berlec, A., and Štrukelj, B. (2020). The Influence of Probiotics on the Firmicutes/Bacteroidetes Ratio in the Treatment of Obesity and Inflammatory Bowel disease. Microorganisms 8. 10.3390/microorganisms8111715.

58. Brook, I. (2007). Treatment of anaerobic infection. Expert Review of Anti-infective Therapy 5, 991–1006. 10.1586/14787210.5.6.991.

59. Rasmussen, M. (2016). Aerococcus: an increasingly acknowledged human pathogen. Clinical microbiology and infection : the official publication of the European Society of Clinical Microbiology and Infectious Diseases 22, 22–27. 10.1016/j.cmi.2015.09.026.

60. Leung, L.H. (1995). Pantothenic acid as a weight-reducing agent: fasting without hunger, weakness and ketosis. Medical hypotheses 44, 403–405. 10.1016/0306-9877(95)90268-6.

61. Suksomboon, N., Poolsup, N., and Darli Ko Ko, H. (2017). Effect of vitamin K supplementation on insulin sensitivity: a meta-analysis. Diabetes, metabolic syndrome and obesity : targets and therapy 10, 169–177. 10.2147/dmso.s137571.

62. Moore, S.A., Hurt, E., Yoder, E., Sprecher, H., and Spector, A.A. (1995). Docosahexaenoic acid synthesis in human skin fibroblasts involves peroxisomal retroconversion of tetracosahexaenoic acid. J Lipid Res 36, 2433–2443.

63. Romano, A., Friuli, M., Eramo, B., Gallelli, C.A., Koczwara, J.B., Azari, E.K., Paquot, A., Arnold, M., Langhans, W., Muccioli, G.G., et al. (2023). "To brain or not to brain": evaluating the possible direct effects of the satiety factor oleoylethanolamide in the central nervous system. Front Endocrinol (Lausanne) 14, 1158287. 10.3389/fendo.2023.1158287.

64. Wang, X., Miyares, R.L., and Ahern, G.P. (2005). Oleoylethanolamide excites vagal sensory neurones induces visceral pain and reduces short-term food intake in mice via capsaicin receptor TRPV1. Journal of Physiology 564, 541–547. 10.1113/jphysiol.2004.081844.

65. Guzmán, M., Lo Verme, J., Fu, J., Oveisi, F., Blázquez, C., and Piomelli, D. (2004). Oleoylethanolamide Stimulates Lipolysis by Activating the Nuclear Receptor Peroxisome Proliferator-activated Receptor α (PPAR-α)*. Journal of Biological Chemistry 279, 27849–27854. 10.1074/jbc.M404087200.

66. Tutunchi, H., Ostadrahimi, A., Saghafi-Asl, M., Hosseinzadeh-Attar, M.-J., Shakeri, A., Asghari-Jafarabadi, M., Roshanravan, N., Farrin, N., Naemi, M., and Hasankhani, M. (2020). Oleoylethanolamide supplementation in obese patients newly diagnosed with non-alcoholic fatty liver disease: Effects on metabolic parameters, anthropometric indices, and expression of PPAR-α, UCP1, and UCP2 genes. Pharmacological Research *156*, 104770. 10.1016/j.phrs.2020.104770.

67. Margheritis, E., Castellani, B., Magotti, P., Peruzzi, S., Romeo, E., Natali, F., Mostarda, S., Gioiello, A., Piomelli, D., and Garau, G. (2016). Bile Acid Recognition by NAPE-PLD. ACS chemical biology 11, 2908–2914. 10.1021/acschembio.6b00624.

68. Pang, Z., Zhou, G., Ewald, J., Chang, L., Hacariz, O., Basu, N., and Xia, J. (2022). Using MetaboAnalyst 5.0 for LC-HRMS spectra processing, multi-omics integration and covariate adjustment of global metabolomics data. Nat Protoc 17, 1735–1761. 10.1038/s41596-022-00710-w.

69. Igarashi, M., Watanabe, K., Tsuduki, T., Kimura, I., and Kubota, N. (2019). NAPE-PLD controls OEA synthesis and fat absorption by regulating lipoprotein synthesis in an in vitro model of intestinal epithelial cells. The FASEB Journal 33, 3167–3179. 10.1096/fj.201801408R.

70. DiPatrizio, N.V. (2016). Endocannabinoids in the Gut. Cannabis and cannabinoid research 1, 67–77. 10.1089%2Fcan.2016.0001.

71. Ridlon, J.M., Kang, D.J., Hylemon, P.B., and Bajaj, J.S. (2014). Bile acids and the gut microbiome. Current opinion in gastroenterology 30, 332–338. 10.1097/mog.0000000000000057.

72. Ramírez-Pérez, O., Cruz-Ramón, V., Chinchilla-López, P., and Méndez-Sánchez, N. (2017). The Role of the Gut Microbiota in Bile Acid Metabolism. Annals of Hepatology 16, S21–S26. 10.5604/01.3001.0010.5672.

73. Hagi, T., Geerlings, S.Y., Nijsse, B., and Belzer, C. (2020). The effect of bile acids on the growth and global gene expression profiles in Akkermansia muciniphila. Applied microbiology and biotechnology 104, 10641–10653. 10.1007/s00253-020-10976-3.

74. Ridlon, J.M., Harris, S.C., Bhowmik, S., Kang, D.-J., and Hylemon, P.B. (2016). Consequences of bile salt biotransformations by intestinal bacteria. Gut microbes 7, 22–39. 10.1080/19490976.2015.1127483.

75. Higuchi, S., Ahmad, T.R., Argueta, D.A., Perez, P.A., Zhao, C., Schwartz, G.J., DiPatrizio, N.V., and Haeusler, R.A. (2020). Bile acid composition regulates GPR119-dependent intestinal lipid sensing and food intake regulation in mice. Gut 69, 1620–1628. 10.1136/gutjnl-2019-319693.

76. Magotti, P., Bauer, I., Igarashi, M., Babagoli, M., Marotta, R., Piomelli, D., and Garau, G. (2015). Structure of human N-acylphosphatidylethanolamine-hydrolyzing phospholipase D: regulation of fatty acid ethanolamide biosynthesis by bile acids. Structure (London, England : 1993) 23, 598–604. 10.1016/j.str.2014.12.018.

77. Proulx, K., Cota, D., Castaneda, T.R., Tschop, M.H., D’Alessio, D.A., Tso, P., Woods, S.C., and Seeley, R.J. (2005). Mechanisms of oleoylethanolamide-induced changes in feeding behavior and motor activity. Am J Physiol Regul Integr Comp Physiol 289, R729–737. 10.1152/ajpregu.00029.2005.

78. Romano, A., Gallelli, C.A., Koczwara, J.B., Braegger, F.E., Vitalone, A., Falchi, M., Micioni Di Bonaventura, M.V., Cifani, C., Cassano, T., Lutz, T.A., and Gaetani, S. (2017). Role of the area postrema in the hypophagic effects of oleoylethanolamide. Pharmacol Res 122, 20–34. 10.1016/j.phrs.2017.05.017.

79. Romano, A., Tempesta, B., Provensi, G., Passani, M.B., and Gaetani, S. (2015). Central mechanisms mediating the hypophagic effects of oleoylethanolamide and N-acylphosphatidylethanolamines: different lipid signals? Front Pharmacol 6, 137. 10.3389/fphar.2015.00137.

80. Rodríguez de Fonseca, F., Navarro, M., Gómez, R., Escuredo, L., Nava, F., Fu, J., Murillo-Rodríguez, E., Giuffrida, A., LoVerme, J., Gaetani, S., et al. (2001). An anorexic lipid mediator regulated by feeding. Nature 414, 209–212. 10.1038/35102582.

81. Fu, J., Gaetani, S., Oveisi, F., Lo Verme, J., Serrano, A., Rodríguez De Fonseca, F., Rosengarth, A., Luecke, H., Di Giacomo, B., Tarzia, G., and Piomelli, D. (2003). Oleylethanolamide regulates feeding and body weight through activation of the nuclear receptor PPAR-alpha. Nature 425, 90–93. 10.1038/nature01921.

82. Wu, F., Guo, X., Zhang, M., Ou, Z., Wu, D., Deng, L., Lu, Z., Zhang, J., Deng, G., Chen, S., et al. (2020). An Akkermansia muciniphila subtype alleviates high-fat diet-induced metabolic disorders and inhibits the neurodegenerative process in mice. Anaerobe 61, 102138. 10.1016/j.anaerobe.2019.102138.

83. Ashrafian, F., Shahriary, A., Behrouzi, A., Moradi, H.R., Keshavarz Azizi Raftar, S., Lari, A., Hadifar, S., Yaghoubfar, R., Ahmadi Badi, S., Khatami, S., et al. (2019). Akkermansia muciniphila-Derived Extracellular Vesicles as a Mucosal Delivery Vector for Amelioration of Obesity in Mice. Front Microbiol 10, 2155. 10.3389/fmicb.2019.02155.

84. Wu, W., Kaicen, W., Bian, X., Yang, L., Ding, S., Li, Y., Li, S., Zhuge, A., and Li, L. (2023). Akkermansia muciniphila alleviates high-fat-diet-related metabolic-associated fatty liver disease by modulating gut microbiota and bile acids. Microb Biotechnol 16, 1924–1939. 10.1111/1751-7915.14293.

85. Depommier, C., Van Hul, M., Everard, A., Delzenne, N.M., De Vos, W.M., and Cani, P.D. (2020). Pasteurized Akkermansia muciniphila increases whole-body energy expenditure and fecal energy excretion in diet-induced obese mice. Gut microbes 11, 1231–1245. 10.1080/19490976.2020.1737307.

86. Keane, J.M., Las Heras, V., Pinheiro, J., FitzGerald, J.A., Nunez-Sanchez, M.A., Hueston, C.M., O’Mahony, L., Cotter, P.D., Hill, C., Melgar, S., and Gahan, C.G.M. (2023). Akkermansia muciniphila reduces susceptibility to Listeria monocytogenes infection in mice fed a high-fat diet. Gut Microbes 15, 2229948. 10.1080/19490976.2023.2229948.

87. Ma, Q., Zhou, X., Su, W., Wang, Q., Yu, G., Tao, W., Dong, Z., Wang, C., Wong, C.M., Liu, T., and Jia, S. (2025). Akkermansia muciniphila inhibits jejunal lipid absorption and regulates jejunal core bacteria. Microbiol Res 293, 128053. 10.1016/j.micres.2025.128053.

88. Matias, I., and Di Marzo, V. (2007). Endocannabinoids and the control of energy balance. Trends Endocrinol Metab 18, 27–37. 10.1016/j.tem.2006.11.006.

89. Everard, A., Belzer, C., Geurts, L., Ouwerkerk, J.P., Druart, C., Bindels, L.B., Guiot, Y., Derrien, M., Muccioli, G.G., Delzenne, N.M., et al. (2013). Cross-talk between Akkermansia muciniphila and intestinal epithelium controls diet-induced obesity. Proceedings of the National Academy of Sciences of the United States of America 110, 9066–9071. 10.1073/pnas.1219451110.

90. Sharma, L., Lone, N.A., Knott, R.M., Hassan, A., and Abdullah, T. (2018). Trigonelline prevents high cholesterol and high fat diet induced hepatic lipid accumulation and lipo-toxicity in C57BL/6J mice, via restoration of hepatic autophagy. Food Chem Toxicol 121, 283–296. 10.1016/j.fct.2018.09.011.

91. Guo, X.F., Sinclair, A.J., Kaur, G., and Li, D. (2018). Differential effects of EPA, DPA and DHA on cardio-metabolic risk factors in high-fat diet fed mice. Prostaglandins Leukot Essent Fatty Acids 136, 47–55. 10.1016/j.plefa.2017.09.011.

92. Zhuge, A., Li, S., Han, S., Yuan, Y., Shen, J., Wu, W., Wang, K., Xia, J., Wang, Q., Gu, Y., et al. (2025). Akkermansia muciniphila-derived acetate activates the hepatic AMPK/SIRT1/PGC-1alpha axis to alleviate ferroptosis in metabolic-associated fatty liver disease. Acta Pharm Sin B 15, 151–167. 10.1016/j.apsb.2024.10.010.

93. Ottman, N., Geerlings, S.Y., Aalvink, S., de Vos, W.M., and Belzer, C. (2017). Action and function of Akkermansia muciniphila in microbiome ecology, health and disease. Best Practice & Research Clinical Gastroenterology 31, 637–642. 10.1016/j.bpg.2017.10.001.

94. Xu, Y., Wang, N., Tan, H.Y., Li, S., Zhang, C., and Feng, Y. (2020). Function of Akkermansia muciniphila in Obesity: Interactions With Lipid Metabolism, Immune Response and Gut Systems. Front Microbiol 11, 219. 10.3389/fmicb.2020.00219.

95. Foley, M.H., O’Flaherty, S., Allen, G., Rivera, A.J., Stewart, A.K., Barrangou, R., and Theriot, C.M. (2021). *Lactobacillus* bile salt hydrolase substrate specificity governs bacterial fitness and host colonization. Proceedings of the National Academy of Sciences 118, e2017709118. 10.1073/pnas.2017709118.

96. Ridlon, J.M., Kang, D.J., and Hylemon, P.B. (2006). Bile salt biotransformations by human intestinal bacteria. J Lipid Res 47, 241–259. 10.1194/jlr.r500013-jlr200.

97. Tang, B., Tang, L., Li, S., Liu, S., He, J., Li, P., Wang, S., Yang, M., Zhang, L., Lei, Y., et al. (2023). Gut microbiota alters host bile acid metabolism to contribute to intrahepatic cholestasis of pregnancy. Nat Commun 14, 1305. 10.1038/s41467-023-36981-4.

98. Repoila, F., Le Bohec, F., Guerin, C., Lacoux, C., Tiwari, S., Jaiswal, A.K., Santana, M.P., Kennedy, S.P., Quinquis, B., Rainteau, D., et al. (2022). Adaptation of the gut pathobiont Enterococcus faecalis to deoxycholate and taurocholate bile acids. Sci Rep 12, 8485. 10.1038/s41598-022-12552-3.

99. Chelakkot, C., Choi, Y., Kim, D.-K., Park, H.T., Ghim, J., Kwon, Y., Jeon, J., Kim, M.-S., Jee, Y.-K., Gho, Y.S., et al. (2018). Akkermansia muciniphila-derived extracellular vesicles influence gut permeability through the regulation of tight junctions. Experimental & molecular medicine 50, e450–e450. 10.1038/emm.2017.282.

100. Guo, S., Nighot, M., Al-Sadi, R., Alhmoud, T., Nighot, P., and Ma, T.Y. (2015). Lipopolysaccharide Regulation of Intestinal Tight Junction Permeability Is Mediated by TLR4 Signal Transduction Pathway Activation of FAK and MyD88. Journal of immunology (Baltimore, Md. : 1950) *195*, 4999–5010. 10.4049/jimmunol.1402598.

101. Wang, F., Graham, W.V., Wang, Y., Witkowski, E.D., Schwarz, B.T., and Turner, J.R. (2005). Interferon-gamma and tumor necrosis factor-alpha synergize to induce intestinal epithelial barrier dysfunction by up-regulating myosin light chain kinase expression. The American journal of pathology 166, 409–419. 10.1016/s0002-9440(10)62264-x.

102. Sidossis, L., and Kajimura, S. (2015). Brown and beige fat in humans: thermogenic adipocytes that control energy and glucose homeostasis. J Clin Invest 125, 478–486. 10.1172/JCI78362.

103. Hamadi, L., and Holliday, J. (2020). Moderators and mediators of outcome in treatments for anorexia nervosa and bulimia nervosa in adolescents: A systematic review of randomized controlled trials. Int J Eat Disord 53, 3–19. 10.1002/eat.23159.

104. Andres-Hernando, A., Jensen, T.J., Kuwabara, M., Orlicky, D.J., Cicerchi, C., Li, N., Roncal-Jimenez, C.A., Garcia, G.E., Ishimoto, T., Maclean, P.S., et al. (2021). Vasopressin mediates fructose-induced metabolic syndrome by activating the V1b receptor. JCI Insight 6. 10.1172/jci.insight.140848.

105. McCormack, S.E., Blevins, J.E., and Lawson, E.A. (2020). Metabolic Effects of Oxytocin. Endocr Rev 41, 121–145. 10.1210/endrev/bnz012.

106. Wahlström, A., Sayin, Sama I., Marschall, H.-U., and Bäckhed, F. (2016). Intestinal Crosstalk between Bile Acids and Microbiota and Its Impact on Host Metabolism. Cell Metabolism 24, 41–50. 10.1016/j.cmet.2016.05.005.

107. Romano, A., Coccurello, R., Giacovazzo, G., Bedse, G., Moles, A., and Gaetani, S. (2014). Oleoylethanolamide: a novel potential pharmacological alternative to cannabinoid antagonists for the control of appetite. Biomed Res Int 2014, 203425. 10.1155/2014/203425.

108. Lauffer, L.M., Iakoubov, R., and Brubaker, P.L. (2009). GPR119 is essential for oleoylethanolamide-induced glucagon-like peptide-1 secretion from the intestinal enteroendocrine L-cell. Diabetes 58, 1058–1066. 10.2337/db08-1237.

109. Caillon, A., Duszka, K., Wahli, W., Rohner-Jeanrenaud, F., and Altirriba, J. (2018). The OEA effect on food intake is independent from the presence of PPARalpha in the intestine and the nodose ganglion, while the impact of OEA on energy expenditure requires the presence of PPARalpha in mice. Metabolism 87, 13–17. 10.1016/j.metabol.2018.06.005.

110. Shea, M.K., Booth, S.L., Gundberg, C.M., Peterson, J.W., Waddell, C., Dawson-Hughes, B., and Saltzman, E. (2010). Adulthood obesity is positively associated with adipose tissue concentrations of vitamin K and inversely associated with circulating indicators of vitamin K status in men and women. J Nutr 140, 1029–1034. 10.3945/jn.109.118380.

111. Zhou, H., Zhang, H., Ye, R., Yan, C., Lin, J., Huang, Y., Jiang, X., Yuan, S., Chen, L., Jiang, R., et al. (2022). Pantothenate protects against obesity via brown adipose tissue activation. Am J Physiol Endocrinol Metab 323, E69–E79. 10.1152/ajpendo.00293.2021.

112. Johansson, M.E.V., Larsson, J.M.H., and Hansson, G.C. (2011). The two mucus layers of colon are organized by the MUC2 mucin, whereas the outer layer is a legislator of host-microbial interactions. Proceedings of the National Academy of Sciences of the United States of America 108 *Suppl*, 4659–4665. 10.1073/pnas.1006451107.

113. van der Lugt, B., van Beek, A.A., Aalvink, S., Meijer, B., Sovran, B., Vermeij, W.P., Brandt, R.M.C., de Vos, W.M., Savelkoul, H.F.J., Steegenga, W.T., and Belzer, C. (2019). Akkermansia muciniphila ameliorates the age-related decline in colonic mucus thickness and attenuates immune activation in accelerated aging Ercc1−/Δ7 mice. Immunity & Ageing 16, 6–6. 10.1186/s12979-019-0145-z.

114. Ottman, N., Reunanen, J., Meijerink, M., Pietilä, T.E., Kainulainen, V., Klievink, J., Huuskonen, L., Aalvink, S., Skurnik, M., Boeren, S., et al. (2017). Pili-like proteins of Akkermansia muciniphila modulate host immune responses and gut barrier function. PloS one 12, e0173004–e0173004. 10.1371/journal.pone.0173004.

115. Gaetani, S., Fu, J., Cassano, T., Dipasquale, P., Romano, A., Righetti, L., Cianci, S., Laconca, L., Giannini, E., Scaccianoce, S., et al. (2010). The fat-induced satiety factor oleoylethanolamide suppresses feeding through central release of oxytocin. Journal of Neuroscience 30, 8096–8101. 10.1523/jneurosci.0036-10.2010.

116. Bucinskaite, V., Tolessa, T., Pedersen, J., Rydqvist, B., Zerihun, L., Holst, J.J., and Hellström, P.M. (2009). Receptor-mediated activation of gastric vagal afferents by glucagon-like peptide-1 in the rat. Neurogastroenterology and motility : the official journal of the European Gastrointestinal Motility Society 21, 978–e978. 10.1111/j.1365-2982.2009.01317.x.

117. Flock, G., Holland, D., Seino, Y., and Drucker, D.J. (2011). GPR119 regulates murine glucose homeostasis through incretin receptor-dependent and independent mechanisms. Endocrinology 152, 374–383. 10.1210/en.2010-1047.

118. Chen, M., Liao, Z., Lu, B., Wang, M., Lin, L., Zhang, S., Li, Y., Liu, D., Liao, Q., and Xie, Z. (2018). Huang-Lian-Jie-Du-Decoction Ameliorates Hyperglycemia and Insulin Resistant in Association With Gut Microbiota Modulation. Front Microbiol 9, 2380. 10.3389/fmicb.2018.02380.

119. Depommier, C., Vitale, R.M., Iannotti, F.A., Silvestri, C., Flamand, N., Druart, C., Everard, A., Pelicaen, R., Maiter, D., Thissen, J.-P., et al. (2021). Beneficial Effects of Akkermansia muciniphila Are Not Associated with Major Changes in the Circulating Endocannabinoidome but Linked to Higher Mono-Palmitoyl-Glycerol Levels as New PPARα Agonists. Cells 10. 10.3390/cells10010185.

