## Supplemental Figures for "Cross-tissue multiomics reveals that *Akkermansia muciniphila* counteracts metabolic syndrome by reprograming gut microbiota, oleoylethanolamide and the gut-hypothalamus axis"

### Figure S1

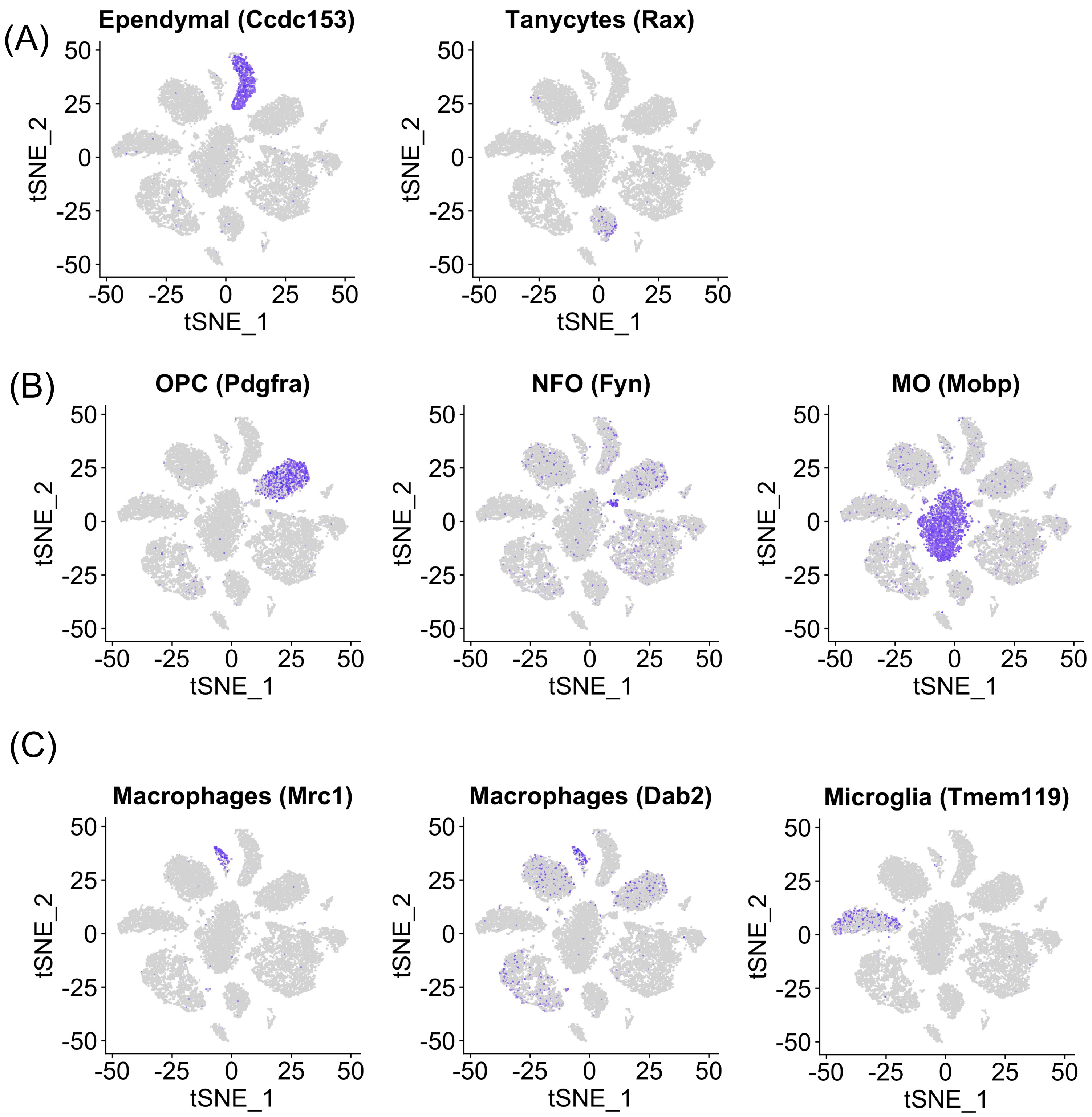
(A)
Ependymal (Ccdc153)
Tanycytes (Rax)
(B)
OPC (Pdgfra)
NFO (Fyn)
MO (Mobp)
(C)
Macrophages (Mrc1)
Macrophages (Dab2)
Microglia (Tmem119)
tSNE_1
tSNE_2
-50
-25
0
25
50

### Figure S2

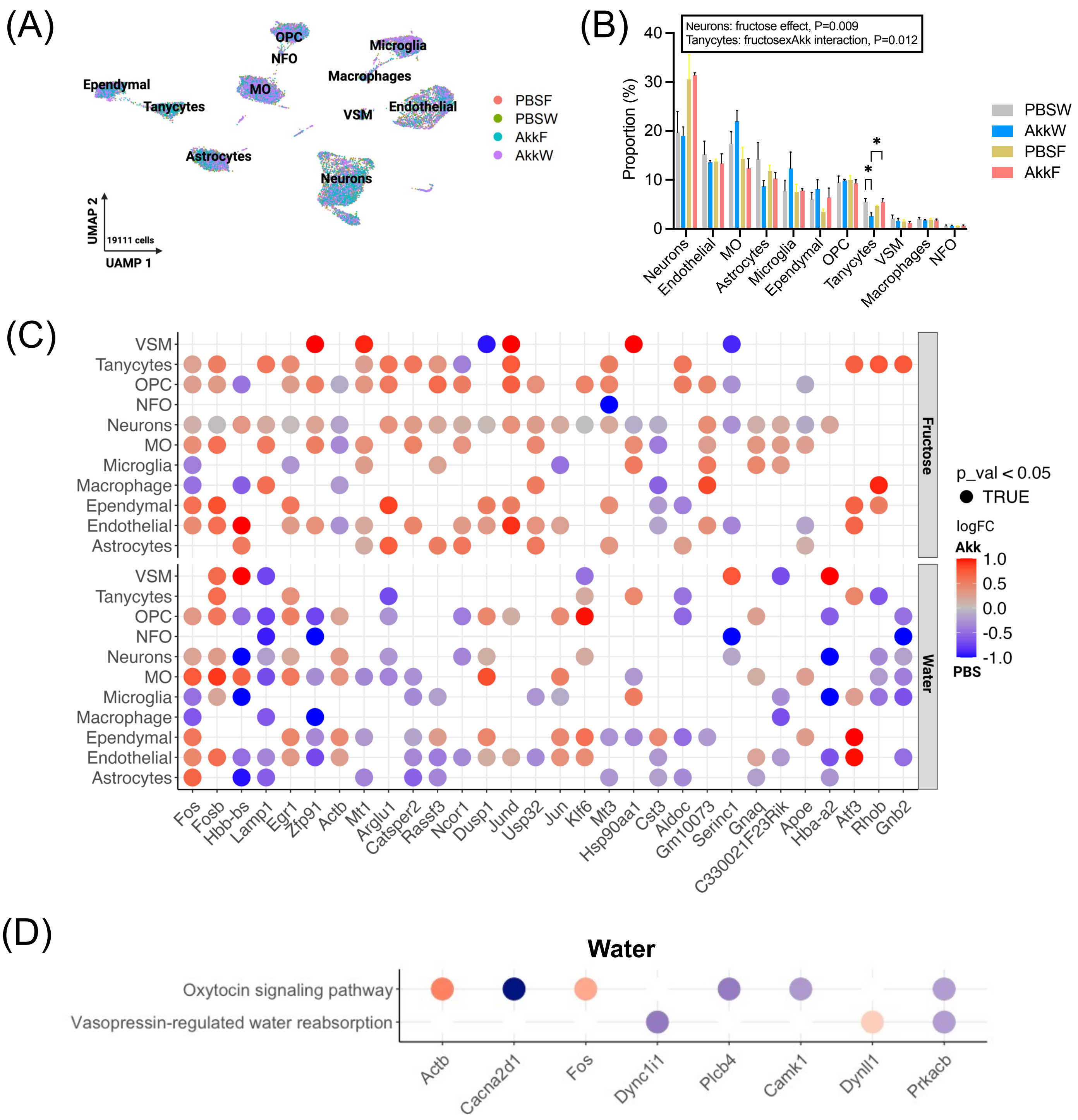
(A)
OPC
NFO
Microglia
Macrophages
Ependymal
MO
Tanycytes
VSM
Endothelial
Astrocytes
Neurons
UMAP 2
19111 cells
UAMP 1
PBSF
PBSW
AkkF
AkkW
(B)
Neurons: fructose effect, P=0.009
Tanycytes: fructosexAkk interaction, P=0.012
Proportion (%)
PBSW
AkkW
PBSF
AkkF
Neurons
Endothelial
MO
Astrocytes
Microglia
Ependymal
OPC
Tanycytes
VSM
Macrophages
NFO
(C)
VSM
Tanycytes
OPC
NFO
Neurons
MO
Microglia
Macrophage
Ependymal
Endothelial
Astrocytes
Fructose
Water
p_val < 0.05
TRUE
logFC
Akk
PBS
Fos
Fosb
Hbb-bs
Lamp1
Egr1
Zfp91
Actb
Mt1
Arglu1
Catsper2
Rassf3
Ncor1
Dusp1
Jund
Usp32
Jun
Klf6
Mt3
Hsp90aa1
Cst3
Aldoc
Gm10073
Serinc1
Gnaq
C330021F23Rik
Apoe
Hba-a2
Atf3
Rhob
Gnb2
(D)
Water
Oxytocin signaling pathway
Vasopressin-regulated water reabsorption
Actb
Cacna2d1
Fos
Dync1i1
Plcb4
Camk1
Dynll1
Prkacb

### Figure S3

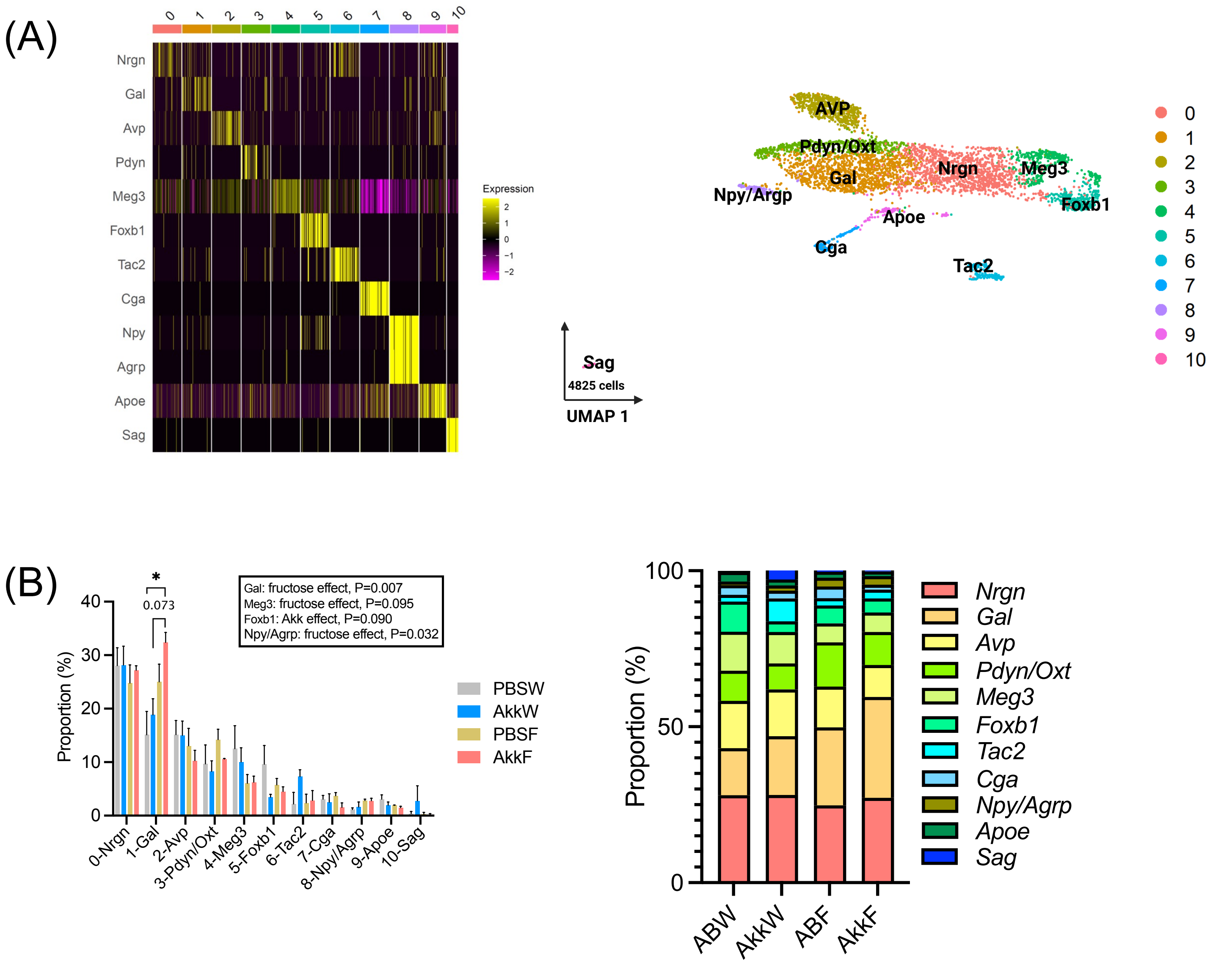
(A)
0 1 2 3 4 5 6 7 8 9 10
Nrgn
Gal
Avp
Pdyn
Meg3
Foxb1
Tac2
Cga
Npy
Agrp
Apoe
Sag
Expression
AVP
Pdyn/Oxt
Gal
Nrgn
Meg3
Foxb1
Npy/Argp
Apoe
Cga
Tac2
Sag
4825 cells
UMAP 1
(B)
Gal: fructose effect, P=0.007
Meg3: fructose effect, P=0.095
Foxb1: Akk effect, P=0.090
Npy/Agrp: fructose effect, P=0.032
0.073
Proportion (%)
PBSW
AkkW
PBSF
AkkF
0-Nrgn
1-Gal
2-Avp
3-Pdyn/Oxt
4-Meg3
5-Foxb1
6-Tac2
7-Cga
8-Npy/Agrp
9-Apoe
10-Sag
Proportion (%)
ABW
AkkW
ABF
AkkF
Nrgn
Gal
Avp
Pdyn/Oxt
Meg3
Foxb1
Tac2
Cga
Npy/Agrp
Apoe
Sag

### Figure S4

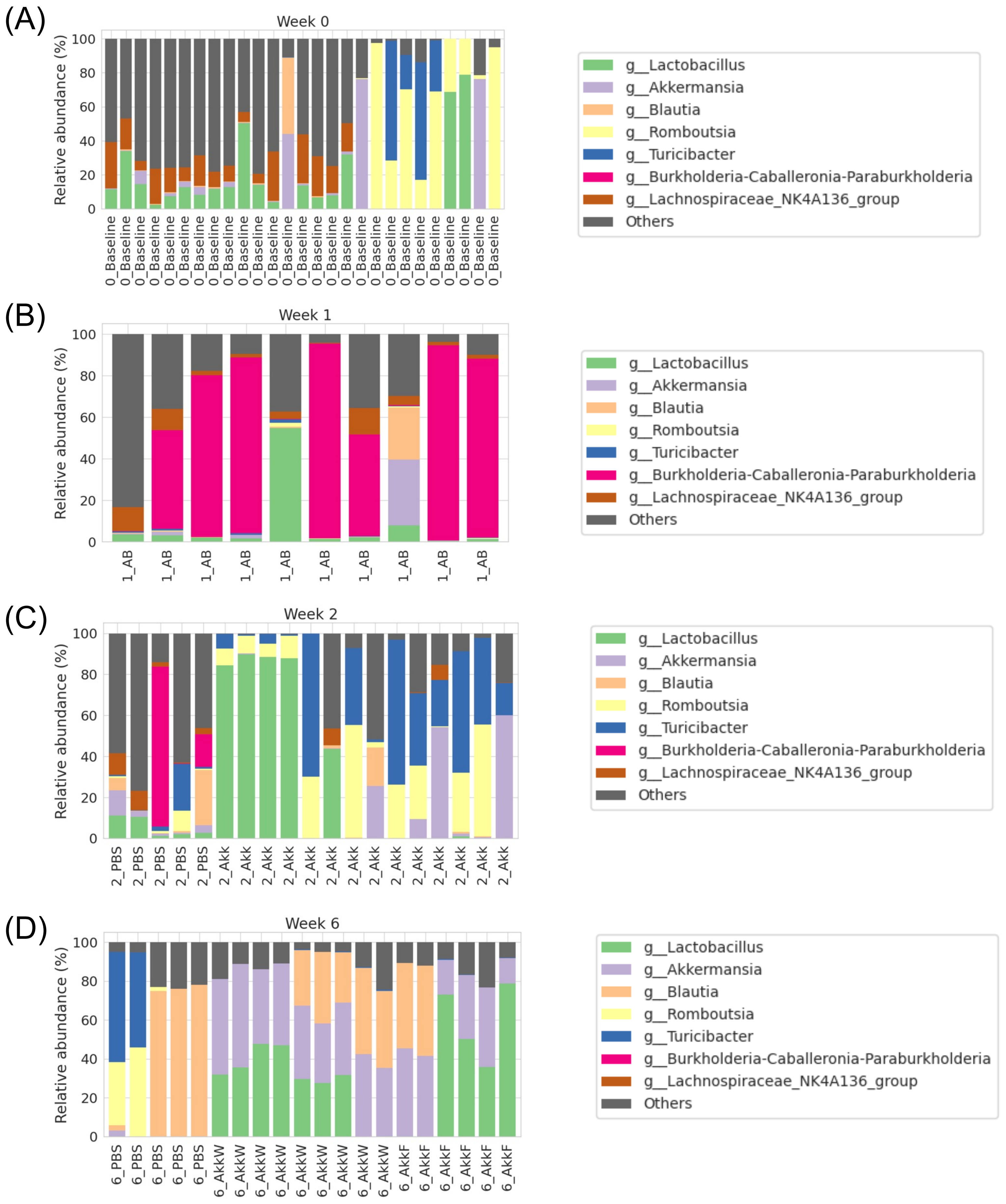
(A)
Week 0
Relative abundance (%)
0
20
40
60
80
100
0_Baseline
g__Lactobacillus
g__Akkermansia
g__Blautia
g__Romboutsia
g__Turicibacter
g__Burkholderia-Caballeronia-Paraburkholderia
g__Lachnospiraceae_NK4A136_group
Others
(B)
Week 1
Relative abundance (%)
1_AB
(C)
Week 2
Relative abundance (%)
2_PBS
2_Akk
(D)
Week 6
Relative abundance (%)
6_PBS
6_AkkW
6_AkkF

### Figure S5

(A)

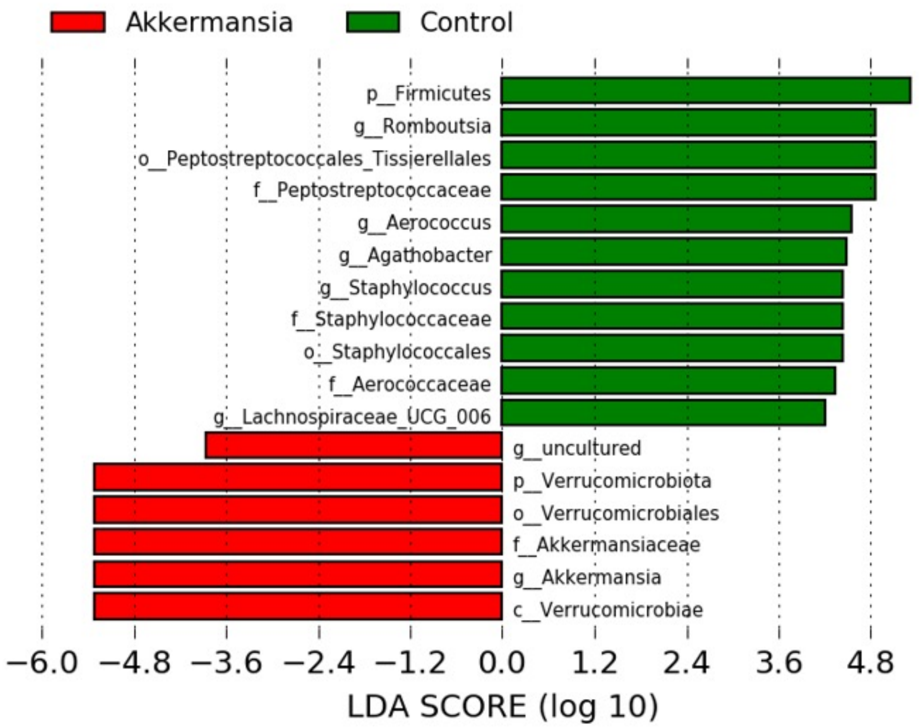

Akkermansia
Control
p__Firmicutes
g__Romboutsia
o__Peptostreptococcales_Tissierellales
f__Peptostreptococcaceae
g__Aerococcus
g__Agathobacter
g__Staphylococcus
f__Staphylococcaceae
o__Staphylococcales
f__Aerococcaceae
g__Lachnospiraceae_UCG_006
g__uncultured
p__Verrucomicrobiota
o__Verrucomicrobiales
f__Akkermansiaceae
g__Akkermansia
c__Verrucomicrobiae
−6.0 −4.8 −3.6 −2.4 −1.2 0.0 1.2 2.4 3.6 4.8
LDA SCORE (log 10)


(B)

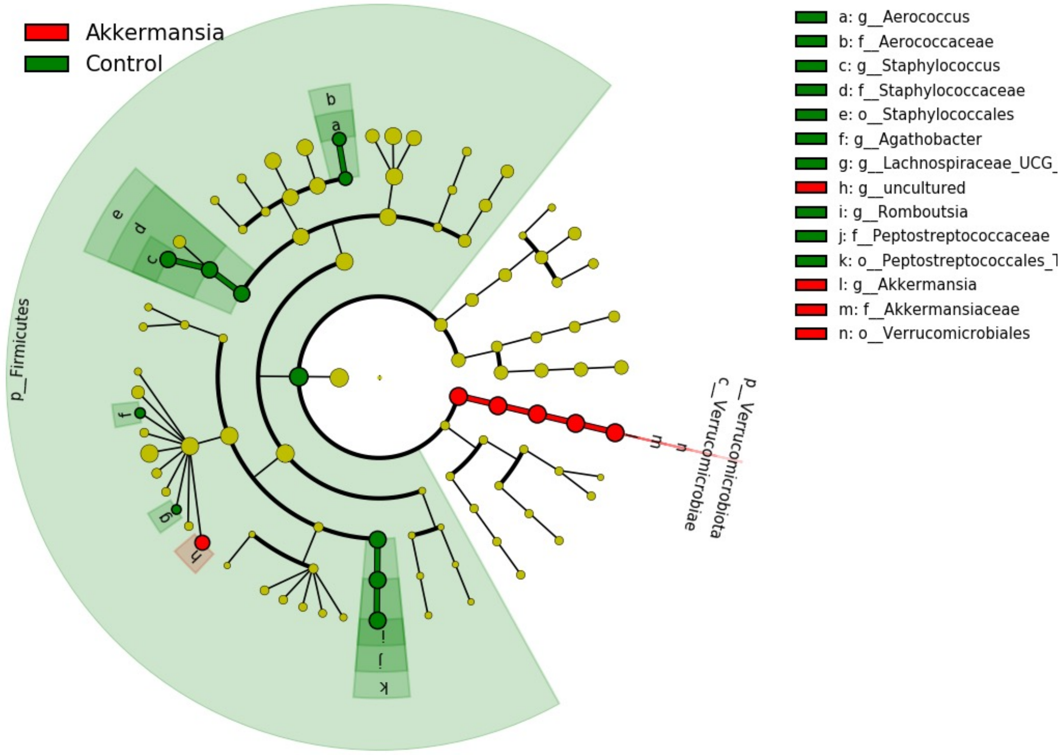

Akkermansia
Control
a: g__Aerococcus
b: f__Aerococcaceae
c: g__Staphylococcus
d: f__Staphylococcaceae
e: o__Staphylococcales
f: g__Agathobacter
g: g__Lachnospiraceae_UCG_
h: g__uncultured
i: g__Romboutsia
j: f__Peptostreptococcaceae
k: o__Peptostreptococcales_T
l: g__Akkermansia
m: f__Akkermansiaceae
n: o__Verrucomicrobiales
p__Firmicutes
p__Verrucomicrobiota
c__Verrucomicrobiae

### Figure S6

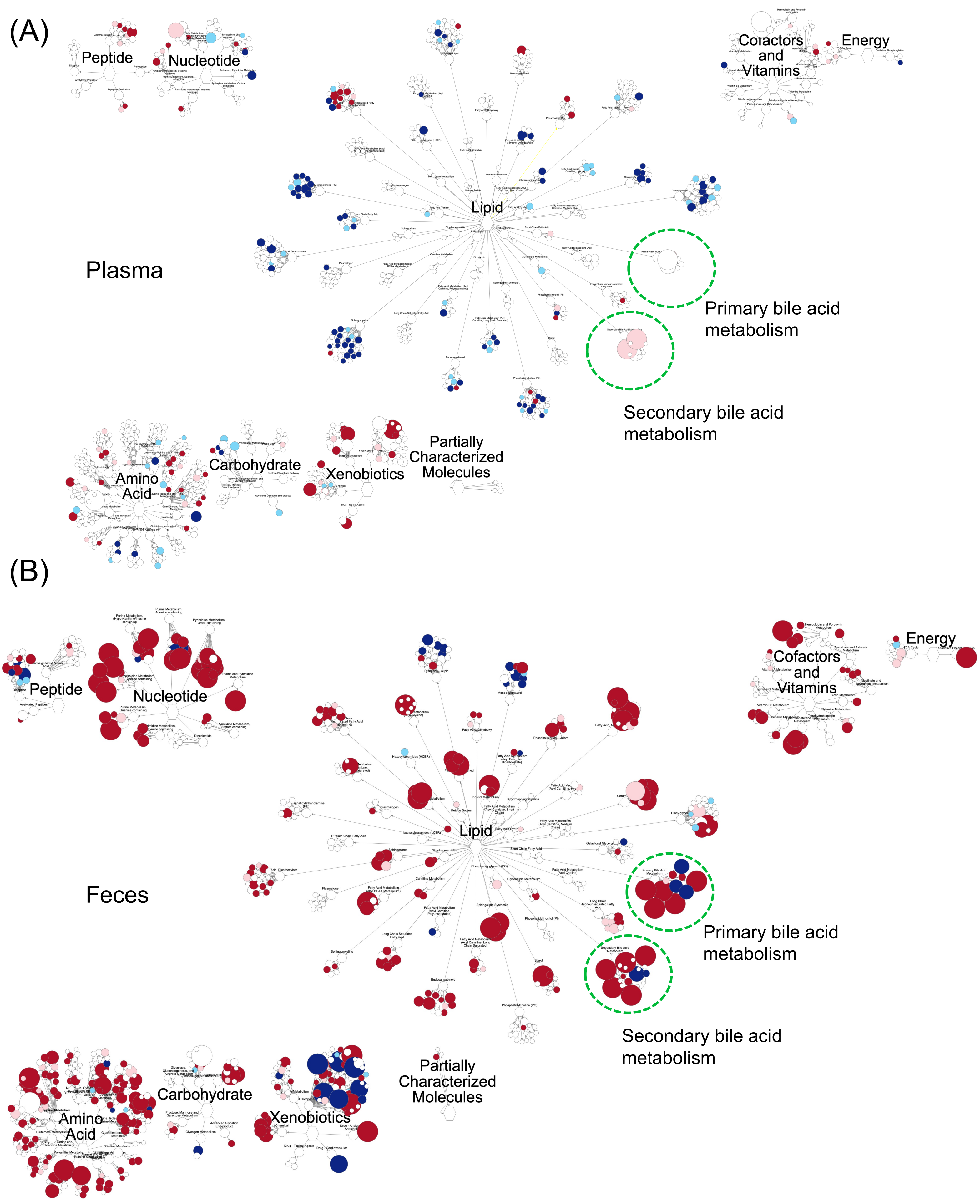
(A)
Peptide
Nucleotide
Lipid
Cofactors and Vitamins
Energy
Plasma
Primary bile acid metabolism
Secondary bile acid metabolism
Amino Acid
Carbohydrate
Xenobiotics
Partially Characterized Molecules
(B)
Peptide
Nucleotide
Lipid
Cofactors and Vitamins
Energy
Feces
Primary bile acid metabolism
Secondary bile acid metabolism
Amino Acid
Carbohydrate
Xenobiotics
Partially Characterized Molecules

### Figure S7

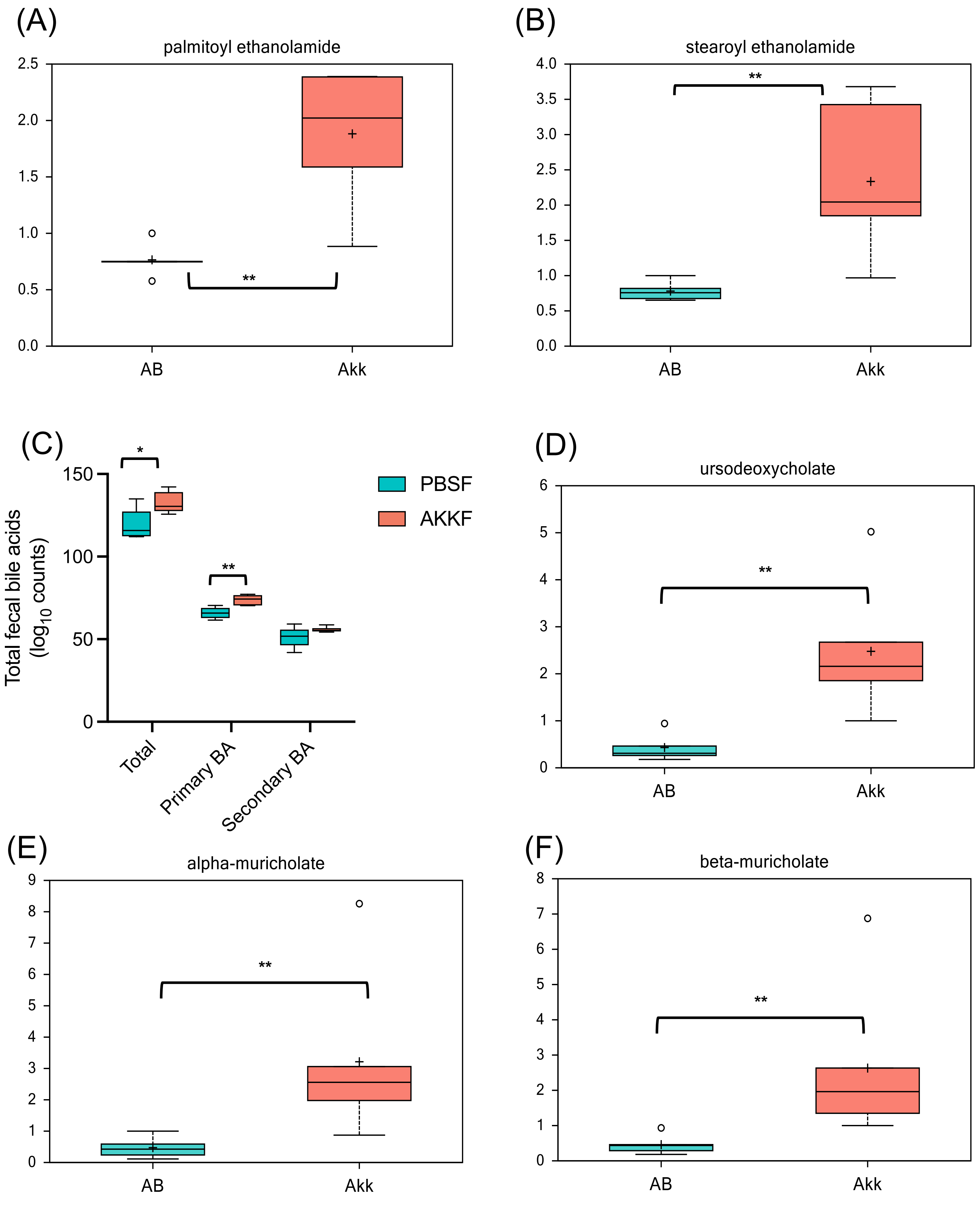
(A)
palmitoyl ethanolamide
2.5
2.0
1.5
1.0
0.5
0.0
**
AB
Akk
(B)
stearoyl ethanolamide
4.0
3.5
3.0
2.5
2.0
1.5
1.0
0.5
0.0
**
AB
Akk
(C)
Total fecal bile acids (log10 counts)
150
100
50
0
*
**
Total
Primary BA
Secondary BA
PBSF
AKKF
(D)
ursodeoxycholate
6
5
4
3
2
1
0
**
AB
Akk
(E)
alpha-muricholate
9
8
7
6
5
4
3
2
1
0
**
AB
Akk
(F)
beta-muricholate
8
7
6
5
4
3
2
1
0
**
AB
Akk

### Figure S8

(A)

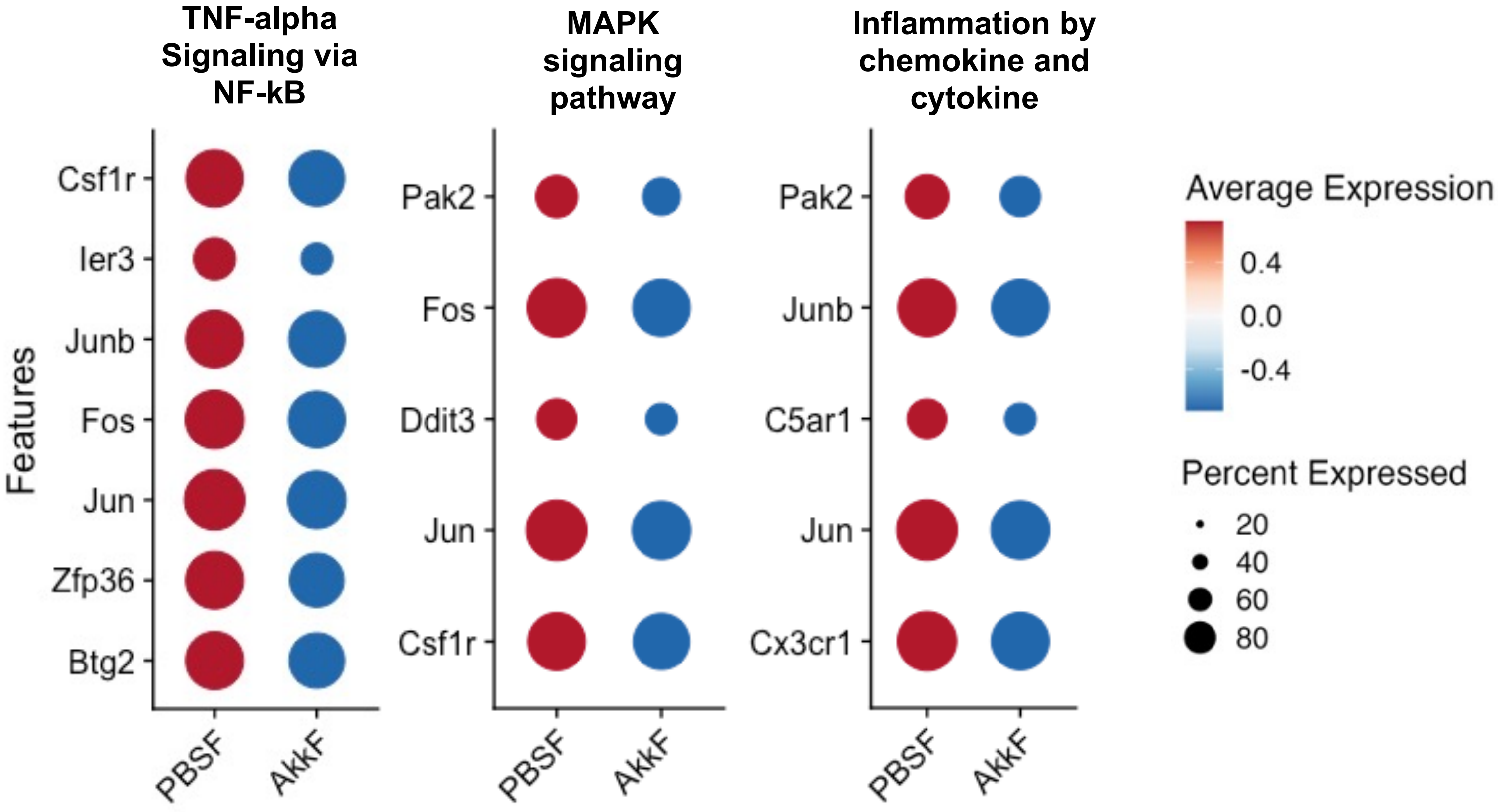
TNF-alpha Signaling via NF-kB
MAPK signaling pathway
Inflammation by chemokine and cytokine
Features
Csf1r
Ier3
Junb
Fos
Jun
Zfp36
Btg2
Pak2
Fos
Ddit3
Jun
Csf1r
Pak2
Junb
C5ar1
Jun
Cx3cr1
PBSF
AkkF
Average Expression
0.4
0.0
-0.4
Percent Expressed
20
40
60
80

### Supplemental Figure legends

**Figure S1. Ependymal, oligodendrocyte and microglia were further differentiated into subtypes based on the marker gene expression.** (A) Ependymal cells were divided into ependymal (*Ccdc153*) and tanycytes based on the expression of tanycyte specific genes (*Rax*). (B) Oligodendrocytes were separated into oligodendrocytes precursor cells (OPC; top marker *Pdgfra*), newly formed oligodendrocytes (NFO; top marker *Fyn*), and myelinating oligodendrocyte (MO; top marker *Mobp*). (C) Macrophages were determined based on the expression of *Mrc1* and *Dab2* gene. A microglia-specific marker gene (*Tmem119*) was used to identify microglia cells. All markers were obtained from Chen et al [35], Romanov et al [34], Hammond et al [36], and Bennett et al [50].

**Figure S2. The effect of *A. muciniphila* on hypothalamus in scRNA-seq.** (A) UMAP plot showing 11 cell types. (B) Cell type proportion. The P values of main factors (fructose, Akk) and interaction were determined by Two-way ANOVA with Tukey's post hoc test in each cell type. Sample size n = 3 per group. * p-val < 0.05. (C) The top 30 most altered DEGs by A. muciniphila across 11 cell types under fructose or water diet, that are ranked by logFC (fold change). (D) Dot plots depicting differentially regulated genes in oxytocin signaling pathway and vasopressin-regulated water reabsorption enriched in neuronal cell type under water condition.

**Figure S3. The effect of *A. muciniphila* on neuronal subtypes in scRNA-seq.** (A) Heatmap showing expression of marker gene that are uniquely expressed in neuronal subclusters and tSNE plot showing a total of 11 subtypes of neurons. (B) Neuronal subtype proportion. Proportions were analyzed using Two-way ANOVA to access the main effects of fructose and Akk and their interaction, followed by Tukey's post hoc test within each cell type. Sample size n = 3 per group. * p-val<0.05.

**Figure S4. Overall microbiome taxonomic profiles depicting changes in microbiome community structure.** Taxa bar plots of fecal microbiota at (A) Week 0 (baseline). (B) Week 1 (after 1 week of antibiotic treatment). (C) Week 2 (after 1 week of *A. muciniphila* treatment). (D) Week 6 (after 5 weeks of *A. muciniphila* treatment and 4 weeks of fructose treatment).

**Figure S5. LefSe results in a bar plot and a cladogram format.** (A) Linear discriminant analysis (LDA) effect size (LEfSe) was used to identify taxa that discriminated between Akk and control (PBS) group at week 6 using standard parameters (p < 0.05, LDA score > 2.0). (B) Cladogram shows the significant taxa at all taxa levels (colored circles). Colors indicate the mouse group (red, green for Akkermansia, Control, respectively) with the highest mean of differential features for which significant differences between groups were found. Taxa (yellow) that do not show significant difference in abundance between groups are unlabeled. n = 5-9/group.

### Supplemental Figure legends

**Figure S6. Network representation of metabolites by known pathways.** OEA is consistently upregulated (red) in both plasma (top) and fecal (bottom) samples. Overall bile acid pool sizes were increased in fecal samples but not in plasma samples. Metabolites of all 9 pathways altered by A. muciniphila under fructose in (A) plasma and (B) feces. Red and blue dots indicate metabolites increased and decreased at p-val < 0.05, respectively, in A. muciniphila-treated mice compared with controls under fructose. n = 5-6/group.

**Figure S7. Box plot showing significantly increased fecal metabolites in the N-acylethanolamine (NAE) family and bile acids upon A. muciniphila treatment.** Box plot showing the levels of fecal (A) Palmittoyl ethanolmide (B) stearoyl ethanolamide, (C) total bile acid pool size (D) ursodeoxycholate (E) alpha-muricholate and (F) beta-muricholate in the PBSF and AkkF groups. Levels are significantly higher in the Akk group. All these metabolites are directly or indirectly related to oleoylethanolamide (OEA) synthesis, suggesting a potential mechanism by which A. muciniphila modulates metabolic processes through NAE family and bile acid pathways. Statistical significance was determined by Welch's t-Test (*q-val < 0.05, **q-val < 0.01)

**Figure S8. A. muciniphila is associated with downregulation of genes involved in inflammatory signaling pathways.** (A) Dot plots showing the expression of genes associated with inflammatory pathways in microglia cells. Under fructose (PBSF) conditions, we observed relatively strong inflammatory signals, including genes in TNF-α signaling via NF-κB, MAPK signaling, and chemokine/cytokine pathways. In contrast, these signals were markedly reduced in the AkkF group, indicating that A. muciniphila supplementation alleviates fructose-induced inflammation.
